# Chemoproteomics discovery of a CNS-penetrant covalent inhibitor of PIKfyve

**DOI:** 10.64898/2026.01.26.701341

**Authors:** Antony J. Burton, Louis S. Chupak, Alison J. Davis, Ahmed S. A. Mady, Mirco Meniconi, Barry Teobald, Bryan W. Dorsey, Lauren R. Byrne, Ryan Mulhern, Berent Lundeen, Elizabeth W. Sorensen, Bharti Patel, Sean Brennan, Dhiraj Kormocha, Ruben Tommasi, Graham L. Simpson, Jeffrey W. Keillor, Laura D’Agostino, Pearl S. Huang, Elayne Penebre

## Abstract

PIKfyve is a lipid kinase involved in regulating protein clearance mechanisms and is a promising target for the treatment of neurodegenerative diseases. Here, we present the discovery and optimization of a CNS-penetrant covalent PIKfyve inhibitor, DUN’058, which achieves sustained target occupancy *in vivo*. Covalent screening hits, identified from chemoproteomics experiments performed in live cells, were rapidly optimized to deliver a brain-penetrant covalent inhibitor of PIKfyve. This covalency centered approach employed a suite of mass spectrometry, biochemical and *in vivo* assays to optimize compound potency, selectivity, and CNS permeability. The target nucleophile, cysteine 1970, is on a flexible loop that appears distal from the kinase active site, highlighting the power of chemoproteomics screening to identify novel nucleophilic amino acids for covalent modification. DUN’058 achieves efficient covalency at the target cysteine, as well as highly selective covalent and reversible selectivity profiles. Covalent PIKfyve inhibition results in modulation of downstream pathway activity, including activation of the transcription factor TFEB, upregulation of protein clearance pathways, and increased GPNMB transcription and secretion of exosome markers. When dosed *in vivo*, DUN’058 achieves sustained target occupancy in the brains of mice long after systemic compound clearance, holding promise for achieving a sustained duration of action in the CNS at low doses, without prolonged effects in the periphery. Taken together, the development of DUN’058 is a demonstration of chemoproteomics-based discovery for a high value CNS target, providing an orally bioavailable and covalent PIKfyve inhibitor.

## 1. Introduction

Neurodegenerative diseases lead to the loss of function and eventual death of nerve cells in the brain and continue to affect an increasing number of people worldwide.^1^ Pathological protein aggregates are hallmarks of neurodegenerative disorders including amyotrophic lateral sclerosis (ALS), frontotemporal dementia (FTD), and Parkinson’s disease.^2-4^ Exocytosis, autophagic and endolysosomal pathways are critical for the removal of such aggregates, presenting pharmacological modulation of proteins associated with such pathways as promising opportunities for neurodegenerative disease amelioration.

PIKfyve is a lipid kinase, and the sole kinase known to phosphorylate phosphatidylinositol-3-phosphate (PI3P) to generate phosphatidylinositol 3,5-bisphosphate (PI(3,5)P_2_).^5, 6^ PIKfyve plays a key role in the endolysosomal system by regulating membrane trafficking, lysosomal biogenesis and autophagy.^7-11^ This occurs through the regulation of PI(3,5)P_2_ and the transcription factor EB (TFEB),^12^ both serving as key signalling molecules. Recent data shows that inhibition of PIKfyve leads to a reduction in pathological aggregates and resultant protection from neuronal toxicity and cell death.^13-16^ Therefore, PIKfyve inhibitors offer strong therapeutic potential for the treatment of neurodegenerative diseases. Indeed, mouse genetic studies have shown that a 50% loss of PIKfyve (through both heterozygous knockout and antisense oligonucleotide (ASO) treatment) results in improved clinical phenotypes and increased survival.^13, 17^ Importantly, it is also known that complete knockout of PIKfyve is not tolerated in mice,^18^ highlighting the importance of achieving precise levels of inhibition both in the brain and the periphery.

Several reversible inhibitors of PIKfyve have been developed and have shown promise in pre-clinical efficacy models, including improvements in clinical signs and modest clearance of protein aggregates in neurons.^19, 20^ We hypothesized that a CNS-penetrant covalent PIKfyve inhibitor would achieve prolonged target engagement in the brain compared to the periphery based on slower protein resynthesis rates in the CNS.^21^ In addition, decoupling of pharmacokinetics and target engagement allows for prolonged target occupancy after the compound has been cleared from circulation, thus lowering the total drug burden and potential for both on-target, off-tissue, as well as off-target liabilities.^22-25^ Taken together, a low-dose, oral, CNS-penetrant, covalent PIKfyve inhibitor has the potential to improve outcomes for patients with neurodegenerative diseases, including ALS, FTD and Parkinson’s disease.

Here, we report the chemoproteomics-based discovery and optimization of a first-in-class CNS-penetrant covalent inhibitor of PIKfyve, capable of achieving quantifiable and sustained CNS target occupancy *in vivo*.

## 2. Results

### 2.1 Discovery of covalent PIKfyve ligands by chemoproteomics screening

As no covalent inhibitors of PIKfyve have been reported, we turned to a cellular chemoproteomics screening approach for hit identification from our covalent compound library. This unbiased activity-based protein profiling (ABPP) approach enables proteome-wide assessment of covalent compound reactivity against native cysteine (Cys) residues across a range of cell types.^26-29^ Cells were treated with compound, lysed, and a desthiobiotin-iodoacetamide (DTB-IA) probe was added to react with unliganded, reactive and accessible Cys residues. The samples were digested and the DTB-labeled peptides were enriched by streptavidin pulldown. Covalent Cys occupancy was inferred from depletion of Cys peptides relative to DMSO control samples, as determined by proteome-wide mass spectrometry (MS; Supplementary methods). This workflow routinely quantifies >20,000 Cys-containing peptides by data-dependent (tandem mass tag-based) and data-independent acquisition (DIA) workflows.^30, 31^ Comparison of DMSO-versus compound-treated cells returns a competition ratio (CR) across the quantified Cys residues, allowing compound- and target-centric evaluation of potentially actionable Cys residues (Fig. 1a).

**Figure 1.**
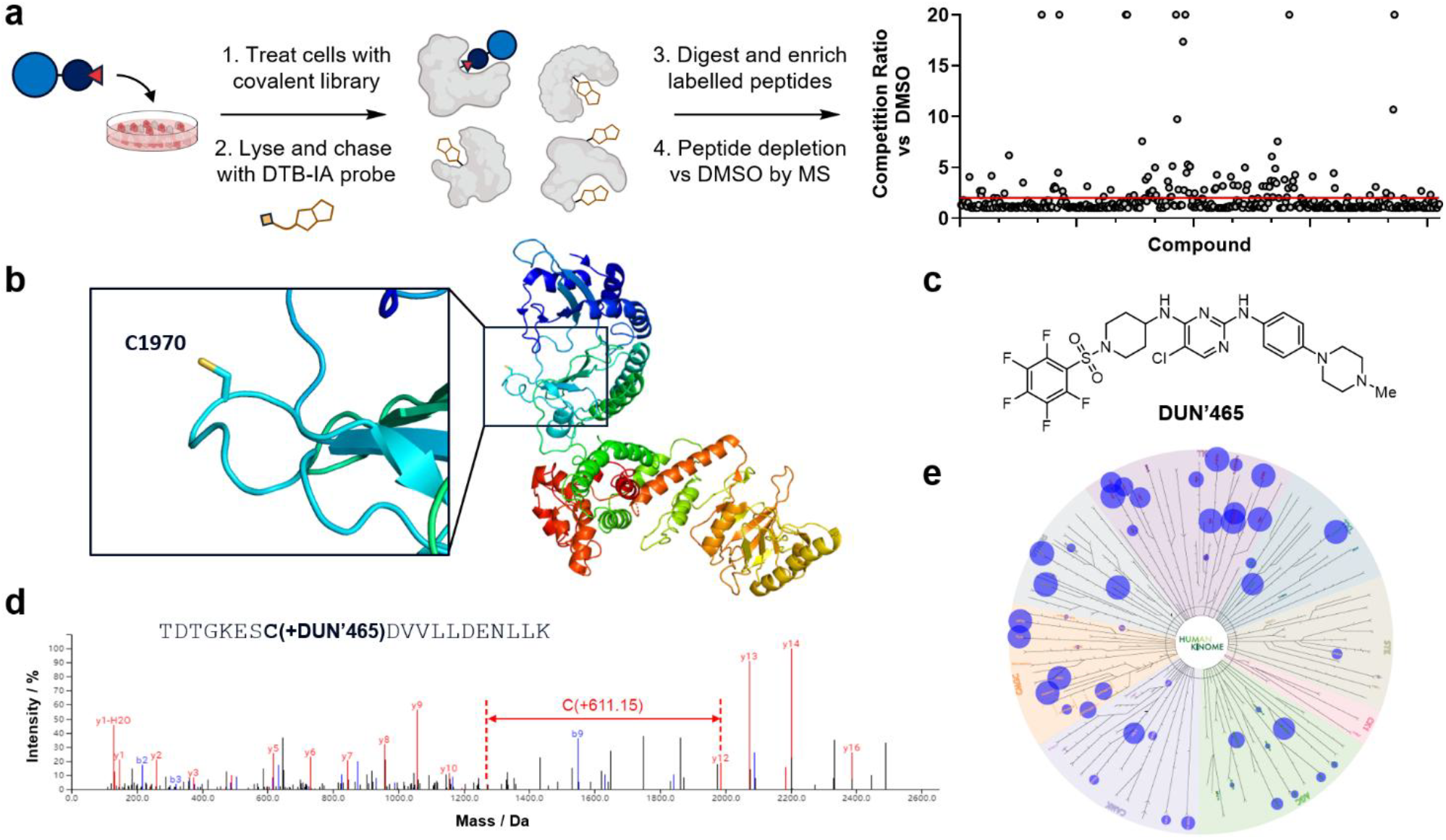
Chemoproteomics screening identifies a ligandable cysteine on the lipid kinase PIKfyve. **a:** Left: Schematic for the ABPP-based chemoproteomics screening workflow. Cells are treated with 10 or 50 µM compound for 2 h, lysed, and treated with a desthiobiotin-iodoacetamide (DTB-IA) probe to label reactive and accessible Cys residues. Peptides are analysed by MS, with a competition ratio (CR) derived from the peptide intensity in DMSO-treated versus compound-treated cells. Right: Chart displaying the CR values for the peptide containing PIKfyve C1970 across the screening library. The hit threshold is set at ≥ 50% target occupancy (CR≥ 2). **b:** PIKfyve domain architecture, highlighting C1970 in an unstructured loop of the kinase domain as resolved by cryo-EM (PDB 7k2v). **c:** Chemical structure of hit compound DUN’465. **d:** Peptide mapping with recombinant, full-length PIKfyve confirms covalency at C1970 (DUN’465 MW = 632.1 Da, adduct MW = 611.2 Da). **e:** Wild-type kinase selectivity profile for DUN’465. The compound inhibits 27 wild-type kinases with ≥ 50% inhibition. The experiment was performed with 66 wild-type and 4 lipid kinases at 1 µM compound in the presence of 10 µM ATP with 2 h incubation at room temperature. The figure was generated using the Reaction Biology Corporation Kinase Mapper.

As part of this hit generation workflow, multiple hits were observed against a previously unliganded Cys on PIKfyve (C1970; Fig. 1a). PIKfyve C1970 is resolved in an unstructured loop of the kinase domain in the only published cryo-EM structure,^32^ and by AlphaFold predictions (Fig. 1b). In total, we observed 8 library compounds that returned a CR of > 20 (> 95% target occupancy (TO)) and 31 compounds with a CR > 4 (> 75% TO) from ABPP experiments conducted in HEK 293T cells with 2 h of compound treatment (SI Table 1). One compound, DUN’465, returned > 90% TO at PIKfyve C1970 with 10 µM compound treatment, with few covalent off targets observed by ABPP (Fig. 1c; SI Fig. 1; SI Table 2). We then confirmed the direct covalency of DUN’465 to C1970 by peptide mapping, observing fragment ions consistent with a single nucleophilic aromatic substitution at the pentafluorobenzenesulfonamide (PFBS; DUN’465 MW = 632.1 Da, observed adduct = 611.2 Da; Fig. 1d; SI Fig. 2). Despite promising covalent selectivity, DUN’465 had suboptimal kinase selectivity. Of 70 kinases evaluated, 27 were inhibited > 50% at 1 µM (Fig. 1e; SI Table 3). Nonetheless, DUN’465 presented multiple avenues for optimization of potency, selectivity and CNS permeability.

### 2.2 Covalency-centred biochemical and biophysical assays enable rapid hit-to-lead discovery

With promising hits obtained from chemoproteomics screening, we developed a strategic suite of assays to understand binding and covalent modification of PIKfyve, thereby guiding hit-to-lead optimization. To this end, full-length (240 kDa), wild-type recombinant PIKfyve (PIKfyve^WT^) was expressed and purified from mammalian cells (SI Fig. 3). Establishing methods to obtain full-length PIKfyve in high purity and good yield was essential, as the isolated kinase domain slowly aggregates over time. An accompanying Cys-Ala (C1970A) mutant full-length protein (PIKfyve^C1970A^) was also expressed and purified to run side-by-side in biochemical and biophysical assays. Importantly, both proteins were active in an ADP-Glo assay, returned similar ATP *K*_M_^app^ values, and were inhibited to equal potency by reversible PIKfyve inhibitors (SI Fig. 4), allowing quantification of the contribution of covalency to PIKfyve inhibition. Potent inhibition of PIKfyve^WT^ was observed after treatment with DUN’465 (IC_50_ (2 h) = 5 nM; Fig. 2a), and a clear potency shift was observed against PIKfyve^C1970A^ (IC_50_ (2 h) > 1,000 nM) consistent with covalency at the target Cys. Next, we sought to establish a screening-scale MS assay to quantify covalency at C1970, improve the assay throughput and decrease the protein requirement of peptide mapping experiments. As full-length PIKfyve (240 kDa) is too large to ionize in routine intact mass analysis (IMA) workflows, we searched for protease cut sites within the protein to selectively liberate the C-terminal kinase domain containing C1970. A single caspase 3 (Casp3) cleavage site was identified after residue 1795 and treatment of recombinant PIKfyve^WT^ with Casp3 resulted in the selective liberation of the kinase domain, as confirmed by LC-MS/MS (Fig. 2b; SI Fig. 5). The digested protein was then treated with compound and analysed by MS, with stable protein signal observed for compound treatments of at least 24 h. Using this assay, we observed covalent addition of DUN’465 to the PIKfyve kinase domain (SI Fig. 6).

**Figure 2.**
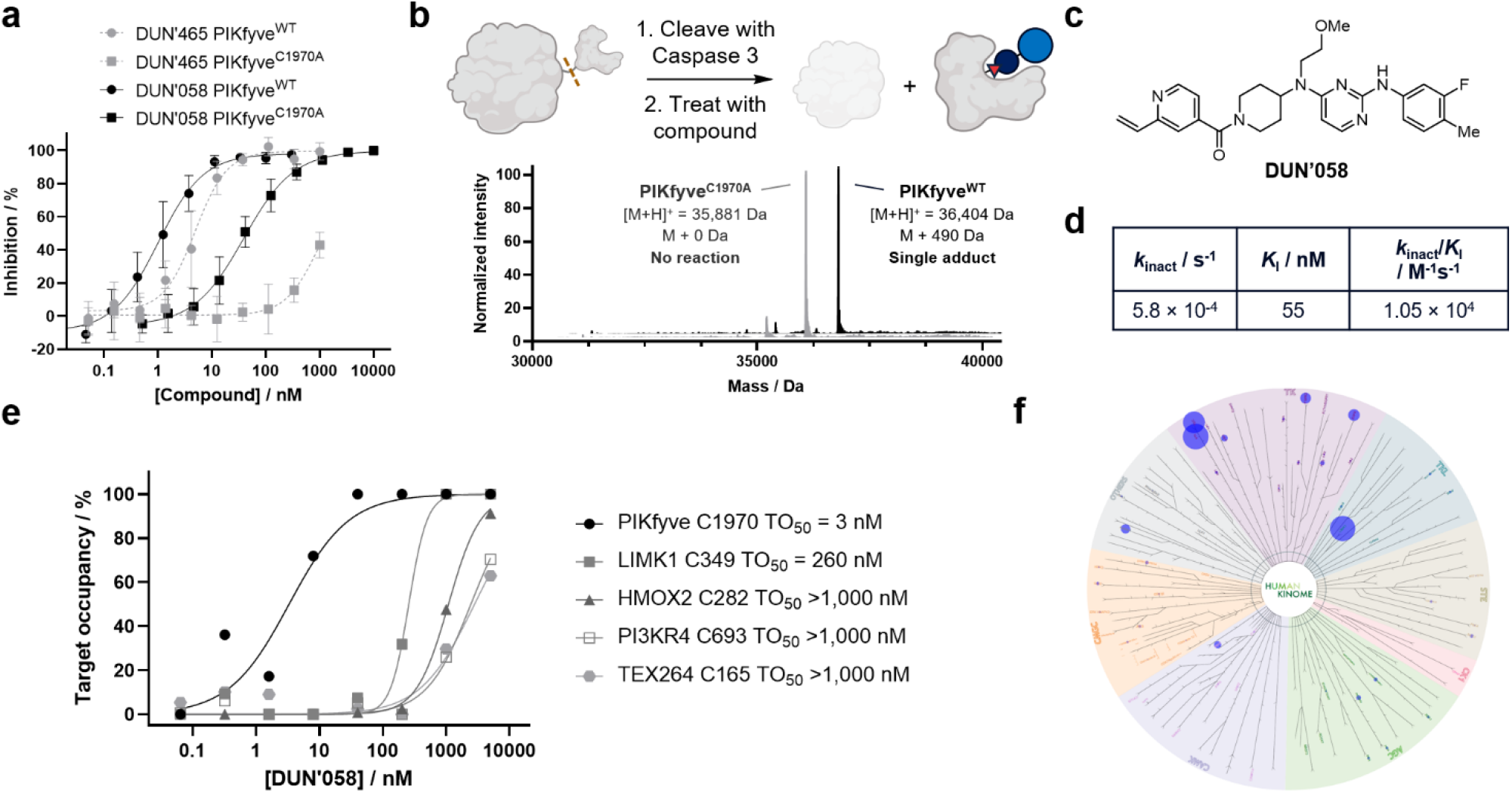
DUN’058 is a covalent and selective inhibitor of PIKfyve C1970. **a:** ADP-Glo assay results for DUN’465 (PIKfyve^WT^ 2 h IC_50_ = 5 nM (grey circles), PIKfyve^C1970A^ 2 h IC_50_ > 1,000 nM (grey squares)) and DUN’058 (PIKfyve^WT^ 2 h IC_50_ < 2.5 nM (black circles), PIKfyve^C1970A^ 2 h IC_50_ = 39 nM (black squares)). A Cys-dependent shift in IC_50_ was observed between PIKfyve^WT^ and PIKfyve^C1970A^, consistent with covalent modification of C1970. **b:** Deconvoluted selective proteolysis-mediated intact mass analysis (IMA) spectra for the reaction of DUN’058 (2.5 µM) with recombinant PIKfyve^WT^ (0.5 µM; black; [M+H]^+^ = 36,404 Da (M + 490 Da) and PIKfyve^C1970A^ (grey; [M+H]^+^ = 35,881 Da (M + 0 Da)). A single adduct was observed between PIKfyve^WT^ and DUN’058 (MW = 490.6 Da) after 2 h incubation at room temperature. **c:** Chemical structure of DUN’058. **d:** *k*_inact_ and *K*_I_ values of DUN’058 measured by time course IMA experiments. **e:** Target occupancy (TO) at PIKfyve C1970 measured in a dose-response ABPP experiment performed in HCT-116 cells, after 2 h treatment with DUN’058. TO_50_ values represent the compound concentration required to achieve 50% TO at the stated time point. Selective covalency at PIKfyve C1970 is observed (TO_50_ = 3 nM). **f:** Wild-type kinase selectivity profile for DUN’058. The compound inhibits 3 wild-type kinases with ≥ 50% inhibition (BTK, TEC, LIMK1). Conditions as in Fig. 1e.

With a bespoke suite of covalency-centred assays established, our optimization campaign focused on potency enhancement, selectivity improvement, oral bioavailability and, critically, CNS penetration. Through systematic analysis, we identified the minimal pharmacophore of DUN’465, conducted comprehensive electrophile scanning to optimize both reversible and covalent binding, and focused on the reduction of molecular weight and polar surface area. These efforts resulted in DUN’058 (Fig. 2c), which exhibits improved physicochemical properties compared to DUN’465, including a reduction in molecular weight (from 632 to 491 gmol^-1^), a decrease in topological polar surface area (from 93 to 82 Å^2^), and a reduction in the number of hydrogen bond donors (from 2 to 1).

DUN’058 achieves exceptional biochemical potency against PIKfyve^WT^ (IC_50_ (2 h) < 2.5 nM in the ADP-Glo assay), with a clear Cys-dependent shift observed with PIKfyve^C1970A^ (IC_50_ (2 h) = 39 nM). The improved potency against PIKfyve^C1970A^ indicates markedly improved reversible binding compared to the chemoproteomics hit. DUN’058 singly modifies the PIKfyve^WT^ kinase domain, and not PIKfyve^C1970A^, in IMA (Fig. 2b) and specifically modifies C1970, as determined by peptide mapping (SI Fig. 7).

Inactivation kinetics are critical parameters to rank covalent efficiency and to guide compound optimization given the time-dependent nature of covalent inhibition.^33-36^ We sought to deconvolute the relative contributions of reversible binding and covalent inactivation for our compounds by independently determining *k*_inact_ and *K*_I_ values. TO for the reaction of DUN’058 with PIKfyve was measured by IMA over a range of concentrations and a 6-hour period. The results were plotted as time versus occupancy across the concentration range tested and *k*_inact_ and *K*_I_ values were then fitted from the measured *k*_obs_ values. DUN’058 exhibited potent reversible binding with *K*_I_ = 55 nM and moderate covalent efficiency with *k*_inact_ = 5.8 × 10^-4^ s^-1^ (Fig. 2d; SI Fig. 8).

DUN’058 maintained the excellent covalent selectivity observed with DUN’465, with only one Cys site beyond PIKfyve C1970, LIMK1 C349, returning a sub-micromolar TO_50_ value, as determined by a 2 h dose-range ABPP experiment (Fig. 2e; SI Table 4). Importantly, we observed dramatically improved kinome selectivity over the screening hit, with > 50% inhibition limited to three kinases after 2 h of treatment with 1 µM DUN’058 (Fig. 2f; SI Table 3). With biochemical, biophysical, and selectivity data in hand, we considered DUN’058 suitable for progression to *in vivo* studies.

### 2.3 C1970 is on a flexible loop and is amenable to covalent modification

As part of our optimization campaign, we developed a non-covalent *in-silico* binding hypothesis for DUN’465 based on the 6.6 Å resolution cryo-EM structure of PIKfyve (PDB 7k2v).^32^ The model provided credible positioning of both the core hinge-binding motif and the peripheral regions of the ligand, in good agreement with crystal structures of similar ligands bound to kinases. In this model, the PFBS electrophile in DUN’465 is located > 12 Å away from C1970 (Fig. 3a). The flexible nature of the loop containing C1970, combined with the low resolution of the cryo-EM structure, suggested it was plausible for the Cys-bearing loop to adopt a conformation consistent with covalent binding.

**Figure 3.**
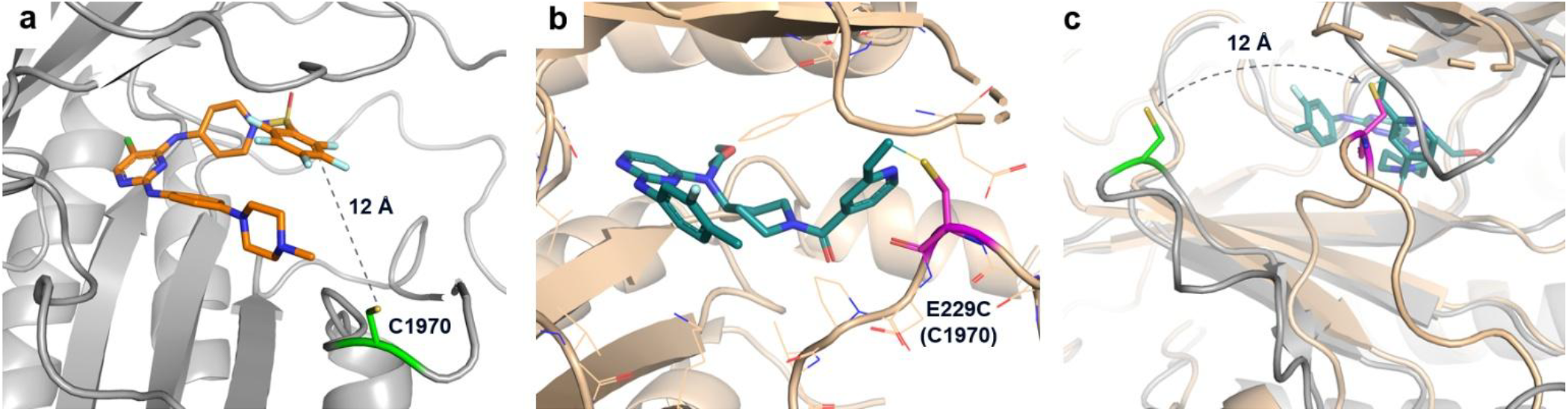
PIP4K2A-PIKfyve chimeric protein confirms binding hypotheses. **a:** *In silico* binding hypothesis of DUN’465 (orange sticks) in the PIKfyve cryo-EM structure (grey ribbons; PDB 7k2v), with C1970 (green sticks) positioned 12 Å from the reactive carbon of DUN’465. **b:** X-ray co-crystal structure of PIP4K2A_6mut_ (tan ribbons) with DUN’058 (teal sticks), with E229C (C1970 in PIKfyve; magenta sticks) positioned for reaction with the vinyl pyridine (PDB: 9zmo). **c**: Overlay of the two structures, showing the flexibility of the C1970-containing loop, facilitating covalency.

We sought to validate our *in-silico* binding hypothesis and covalency at C1970 with X-ray crystallography. As only one structure of full-length PIKfyve has been disclosed and the isolated kinase domain is prone to aggregation during purification, we identified a lipid kinase (PIP4K2A) that is structurally homologous to PIKfyve and amenable to bacterial expression and X-ray crystallography. A chimeric PIP4K2A-PIKfyve protein was designed that incorporated six mutations into the wild-type PIP4K2A sequence (denoted as PIP4K2A^6mut^). These included: 1) 4 mutations to make the PIP4K2A ATP binding site similar in shape and properties to PIKfyve, 2) E229C to mimic C1970 in PIKfyve, and 3) L230D to increase the reactivity of E229C. PIP4K2A^6mut^ was expressed and purified from *E. coli* and, on treatment with DUN’058, showed > 95% single modification by IMA, with modification at C229 confirmed by peptide mapping (SI Figs. 9-11). The purified, adducted PIP4K2A^6mut^ protein was then subjected to crystallization trials. Optimized diffraction-quality crystals resulted in a 3.08 Å resolution co-crystal structure (SI Table 5; PDB: 9zmo). Density for DUN’058 was observed in the kinase active site, and the compound adopted a U-shaped conformation as predicted by computational modelling. Importantly, we observed clear electron density for C229 (C1970 in PIKfyve) positioned adjacent to the vinyl pyridine electrophile, confirming that the loop containing our target Cys is dynamic and able to react with our compounds bound to the kinase active site (Fig. 3b,c). The structure highlights an important advantage of unbiased covalent screening in cells, where an apparently solvent-exposed Cys (as resolved by cryo-EM and predicted by AlphaFold) would not have been considered tractable in the absence of our chemoproteomics hits.

### 2.4 DUN’058 modifies PIKfyve in cells and impacts downstream on-target pathways

Next, we evaluated our PIKfyve inhibitors in live cells to confirm covalent TO and evaluate downstream target engagement events. We established targeted MS-based assays to enable rapid evaluation of covalent TO across a range of biological input materials. Parallel reaction monitoring (PRM)-based MS methods were established to quantify the tryptic peptide containing C1970, along with appropriate control peptides from PIKfyve and four housekeeping proteins (Fig. 4a; Supplementary methods).^37, 38^ HCT-116 cells were treated with a dose range of DUN’058 for 2 h, lysed in denaturing lysis buffer, and processed for MS analysis (Supplementary methods). This depletion-based assay returned a 2 h TO_50_ of 5 nM, aligning well with NanoBRET target engagement assays (Fig. 4b; SI Fig. 12). TO_50_ measurements from mouse brain homogenate spike-in experiments and from human motor neurons (hMN) derived from induced pluripotent stem cells (iPSC) also aligned well with the HCT-116 data, returning TO_50_ values of 6 and 13 nM, respectively, with 1 and 2 h compound incubation times (Fig. 4b). PRM-based targeted MS was also used to detect the direct covalent adduct of DUN’058 to C1970, confirming that the depletion of the C1970 peptide is indeed a result of DUN’058 covalently reacting with PIKfyve.

**Figure 4.**
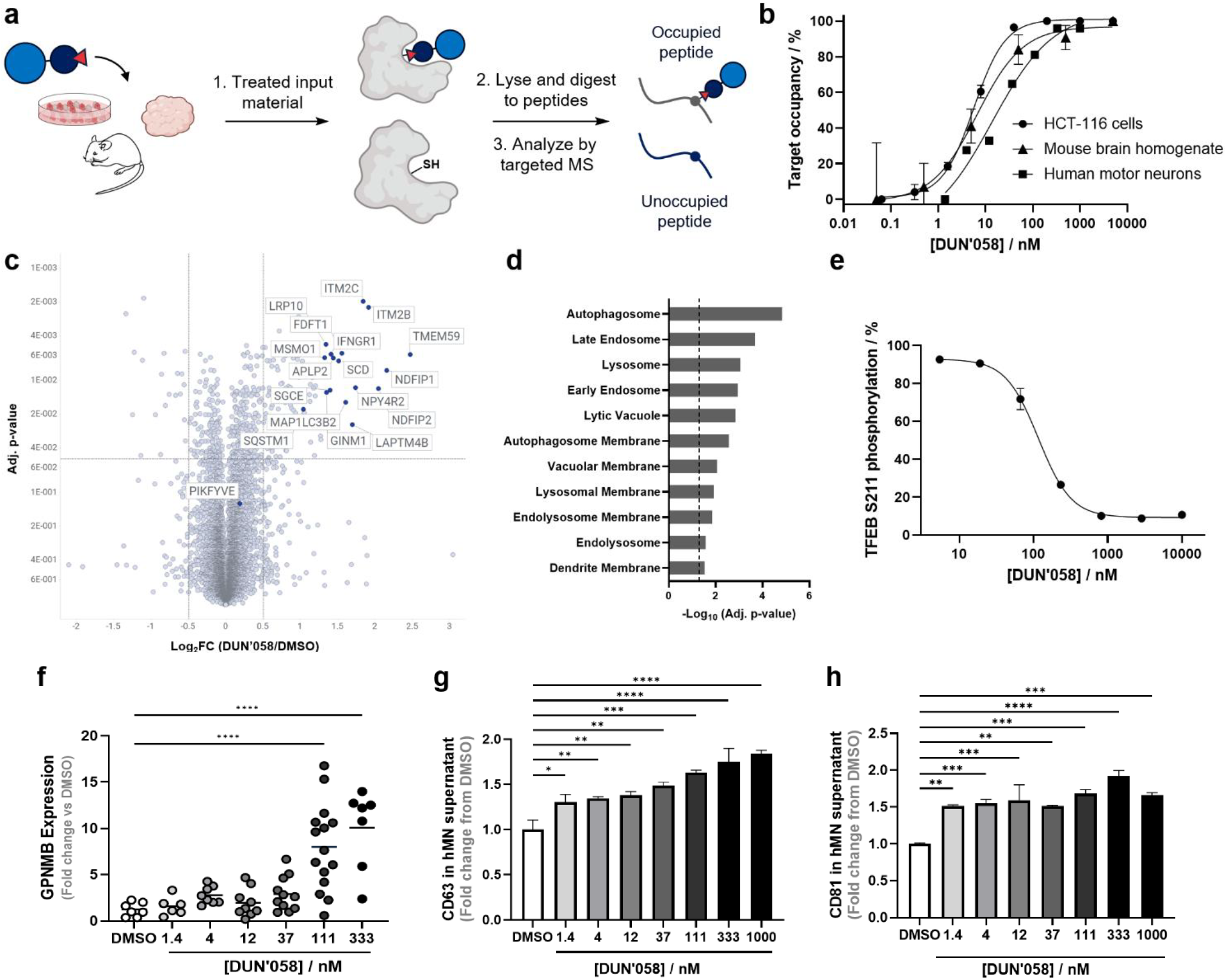
DUN’058 engages C1970 in cells and tissue homogenates and impacts on-target downstream pathways. **a:** Schematic for the PRM-based targeted MS TO assay. **b:** TO_50_ curves for the covalent modification of DUN’058 to PIKfyve C1970 from HCT-116 cells (2 h TO_50_ = 5 nM; circles), mouse brain homogenate (1 h TO_50_ = 6 nM; triangles) and hMN (2 h TO_50_ = 13 nM; squares). **c:** Volcano plot visualizing data from a global proteomics experiment run in HCT-116 cells treated with 500 nM DUN’058 for 24 h. Cutoffs displayed: Log_2_ fold change ≤ -0.5 or ≥ 0.5, adjusted *p*-value < 0.05. **d:** Significantly upregulated cellular component gene ontology terms from the global proteomics experiment in **d**, as identified by Enrichr (cutoff = adjusted *p*-value < 0.05). **e:** Inhibition of TFEB S211 phosphorylation after treatment of SU-DHL-6 cells with DUN’058 for 2 h (IC_50_ = 116 nM). **f:** Quantification of GPNMB mRNA from hMN treated for 24 h with DUN’058, as determined by qPCR. One-way ANOVA (Dunnet), *p*-values, **** < 0.0001. **g**,**h:** Quantification of the exosome markers CD63 (**g**) and CD81 (**h**) in exosomes isolated from the supernatant of hMN treated for 24 h with DUN’058, as determined by Luminex immune capture. One-way ANOVA (Dunnet), *p*-values * < 0.05, ** < 0.01, *** < 0.001, **** < 0.0001.

With cellular TO confirmed, we sought to determine the proteome-wide effects of covalent PIKfyve inhibition in a global proteomics experiment run with DUN’058-treated HCT-116 cells. No change in the abundance of PIKfyve itself, nor two direct interactors of PIKfyve, Fig4 and Vac14,^39^ was observed, as expected with our covalent inhibition mechanism of action. Conversely, proteome changes consistent with selective PIKfyve pathway modulation were clearly observed, including upregulation of proteins associated with endolysosomal pathways and autophagy (*e*.*g*. MAP1LC3B2, SQSTM1; Fig. 4c,d; SI Table 6). Importantly, modulation of off-target pathways or pathways associated with potential safety risks was not seen by global proteomics.

PIKfyve inhibition activates TFEB through perturbation of its interaction with the kinase mTORC1.^12^ This leads to dephosphorylation of TFEB S211 by PP2A, nuclear TFEB translocation, and transcriptional activation of genes required for autophagy and lysosomal function. A decrease in TFEB S211 phosphorylation was indeed observed in SU-DHL-6 cells on treatment with DUN’058, returning an IC_50_ of 116 nM after 2 h of compound treatment, confirming that covalent TO at PIKfyve results in downstream biological effects (Fig. 4e). Next, we evaluated the effects of DUN’058 treatment on a TFEB target gene in the autophagy pathway and an established biomarker of PIKfyve inhibition, the glycoprotein GPNMB.^19^ A dose-dependent increase in GPNMB mRNA was observed by qPCR in cultured iPSC-derived hMN treated with a dose range of DUN’058 for 24 h (Fig. 4f). PIKfyve inhibition is also known to increase exocytosis.^40^ Culture supernatants from hMN, treated with a dose range of DUN’058 for 24 h, were collected and assayed for the presence of CD63- and CD-81 positive exosomes, both of which are used as markers to quantify exosome release in cell culture supernatant.^41^ DUN’058 treatment resulted in a dose-dependent increase in the number of both CD63- and CD81-positive exosomes across a concentration range that aligned well with TO measurements in hMN (Fig. 4g). Taken together, *in vitro* cellular experiments confirmed covalency at our target Cys and downstream on-target pathway effects of covalent PIKfyve inhibition.

### 2.5 DUN’058 is a CNS-penetrant, covalent inhibitor of PIKfyve C1970 *in vivo*

With the comprehensive *in vitro* characterization of DUN’058 complete, we next evaluated our covalent PIKfyve inhibitor in mice to assess the compound’s pharmacokinetics, CNS penetration, *in vivo* TO and safety. A single 50 mg/kg oral dose of DUN’058 was administered to C57BL/6 mice and compound levels were measured in plasma and brain tissue over 4 h. The compound achieved plasma C_max_ levels of 7.8 µM at 0.5 h with concomitant brain levels of 1.8 µM, and a plasma t_1/2_ of 0.6 h (SI Fig. 13). With oral bioavailability and CNS penetration demonstrated, the compound was taken into a mouse study to assess TO in various on- and off-target tissues over 5 days.

A single 150 mg/kg oral dose of DUN’058 was administered to C57BL/6 mice on Day 1 and plasma samples, as well as brain, spleen and duodenum tissues were collected for analysis over 5 days. Analysis of isolated plasma and tissues returned DUN’058 concentrations of 4.8 µM, 0.4 µM and 2.5 µM at 4 h in plasma, brain and spleen, respectively, and the compound was completely cleared by 48 h (Fig. 5a). We next determined PIKfyve TO in brain, spleen and duodenum tissue using our targeted MS assay. A TO_max_ of 80% was achieved in brain tissue at 4 h and TO was observed at time points up to and including 120 h, representing sustained TO well after compound clearance (Fig. 5b; SI Table 7). Dose-proportional TO was observed in the spleen and duodenum, as expected, but with shorter durations of action. We were also able to confirm the presence of the direct covalent adduct of DUN’058 to C1970 in all three tissues, eliminating the possibility that observed TO is the result of an active metabolite (Fig. 5b; SI Table 7).

**Figure 5.**
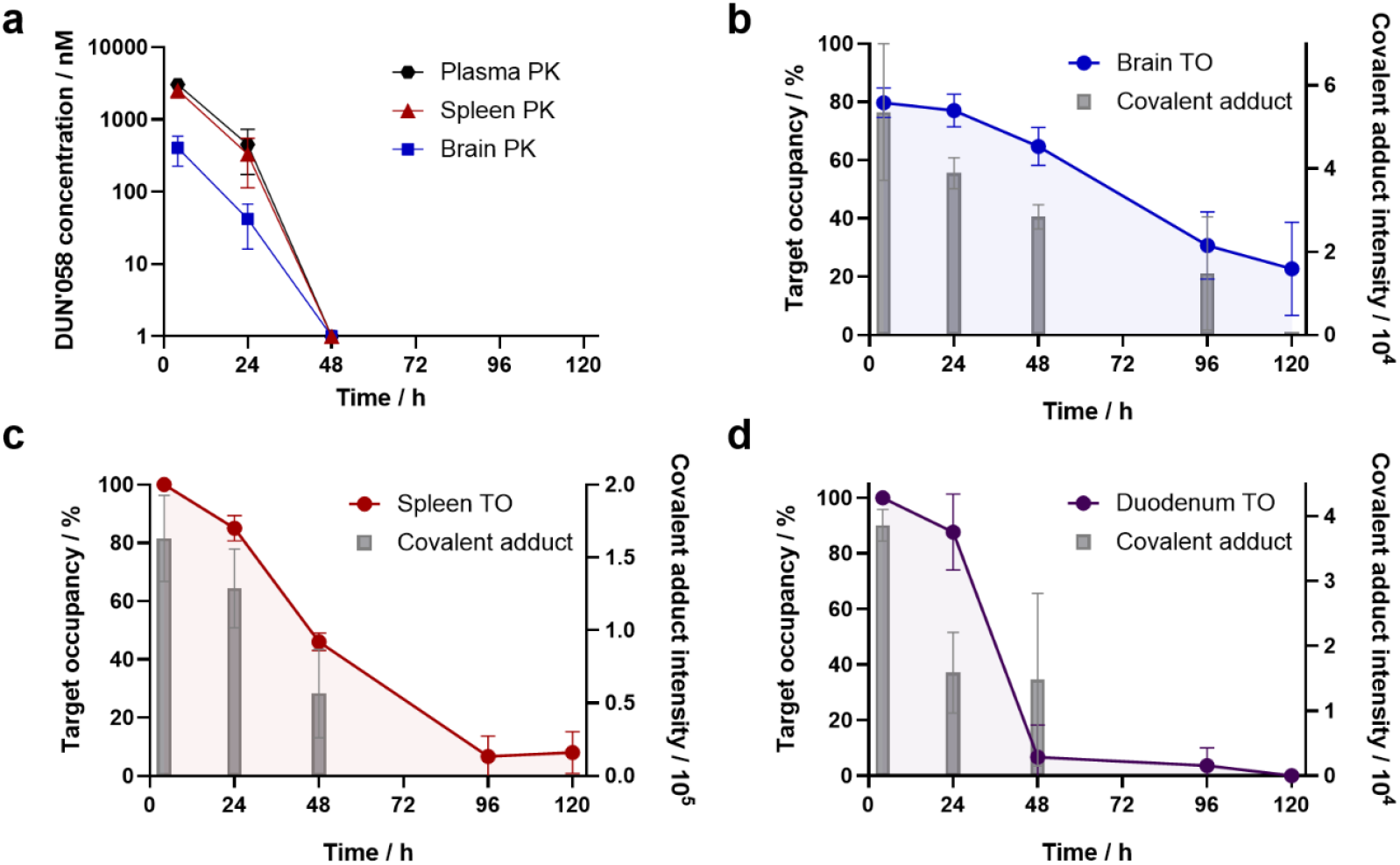
DUN’058 achieves sustained TO in the brains of dosed mice. **a:** Compound exposure in the plasma (black), spleen (red) and brains (blue) of mice after a single 150 mg/kg oral dose of DUN’058. **b:** Sustained TO is observed in the brains of mice dosed as in **a**. Depletion-based TO measurements (blue) align well with detection of the direct covalent adduct of DUN’058 to the C1970-containing peptide (grey). **c&d**: TO determined in the spleen (**c**) and duodenum (**d**) of mice dosed as in **a**. Shorter durations of action are observed in both peripheral tissues when compared with the brain.

With the TO results in hand, we determined the resynthesis rate of PIKfyve in mouse brain by fitting the 5-day TO data to a monoexponential decay model (SI Fig. 14).^42, 43^ Given the irreversible nature of the covalent adduct between DUN’058 and our target Cys, the rate of loss of TO can be directly inferred as the PIKfyve protein resynthesis rate. Conversion of the obtained first-order rate constant to a protein half-life returned a slow PIKfyve resynthesis rate in the brain (t_1/2_ = 72 h). The measured PIKfyve protein resynthesis rate in spleen (t_1/2_ = 35 h) was significantly faster than in brain, and faster still in the duodenum (t_1/2_ < 24 h; SI Fig. 14), consistent with a faster PIKfyve resynthesis rate in peripheral tissues with more rapid cell turnover. Our measured PIKfyve resynthesis rates align well with recent protein half-life values measured in mouse tissues by stable isotope labelling by amino acids in cell culture (SILAC)-based MS proteomics, where PIKfyve has a slower resynthesis rate in mouse brain regions (t_1/2_ ≈ 5 days) when compared with peripheral tissues.^21^

Finally, we performed a 5-day repeat dose study in mice with a daily 50 mg/kg oral dose of DUN’058. Brain TO was measured 0.5 and 24 h after the last dose, with daily dosing achieving sustained PIKfyve TO of ∼85%. Importantly, no weight loss or adverse effects were recorded during the study (SI Fig. 15). Taken together, the *in vivo* data provides a pre-clinical proof-of-concept that a CNS-penetrant covalent PIKfyve inhibitor can achieve durable and safe TO in the brain, alongside reduced durations of engagement in the periphery.

## 3. Discussion

Here, we describe the development of a first-in-class, CNS-penetrant covalent inhibitor of the lipid kinase PIKfyve, discovered by unbiased live-cell chemoproteomics screening. Chemoproteomics screening strategies present opportunities to drug challenging protein targets by identifying and reacting with nucleophilic amino acid side chains, both Cys and beyond, located away from canonical enzyme active sites.^44, 45^ Such covalency events can uncover novel mechanisms of action, including modulation of enzymatic function or by inducing or disrupting protein-protein interactions. Recent studies have highlighted the impact of such strategies for targets including Werner helicase, JAK1, FOXA1, and TRMT112-METTL5.^37, 46-48^ Indeed, the positioning of PIKfyve C1970 observed by cryo-EM and in computational structure prediction would prove challenging to target through rational design, highlighting the power of chemoproteomics-based hit generation to identify non-canonical nucleophiles as starting points for covalent drug discovery.

Our chemoproteomics hits were optimized by covalency-led medicinal chemistry, assembling an efficient assay funnel comprising mass spectrometry-based proteomics, biochemical, biophysical, and cellular biology assays. Several innovations were required to enable this optimization process. A selective-proteolysis mediated IMA assay was developed to rapidly quantify covalent adduct formation for a high molecular weight protein. The relative contributions of *k*_inact_ and *K*_I_ to covalent inactivation were determined by assays deploying full-length and biochemically active PIKfyve^WT^ and PIKfyve^C1970A^ proteins. We designed a novel chimeric protein to obtain a co-crystal structure and confirm our binding hypothesis, thereby facilitating structure-based compound optimization. The resulting compound from these optimization efforts, DUN’058, displays efficient on-target covalent inactivation kinetics together with excellent reversible and covalent selectivity profiles.

When evaluated *in vivo*, DUN’058 achieves sustained TO in the brains of dosed mice with oral administration. Target-tissue selective and durable inhibition arises from slower protein resynthesis rates in the CNS, when compared with peripheral tissues. Moreover, the ability to quantify TO with covalent compounds facilitates the precise optimization of dosing to achieve defined levels of target engagement, representing a level of control that is not easily achieved with reversible compounds. Covalent inhibition of PIKfyve results in the upregulation of protein clearance pathways, highlighting the potential for the use of PIKfyve inhibitors for the treatment of neurodegenerative diseases. Moreover, DUN’058 also holds promise for studying the role of PIKfyve in disease areas beyond neurodegeneration, including cancer and viral infections.^49, 50^

The work described here is a demonstration of the power of chemoproteomics-based hit discovery against a high value CNS target, resulting in an orally bioavailable and covalent PIKfyve inhibitor amenable to precision dosing and determination of target occupancy, both in the brain and periphery.

## Acknowledgements

We thank WuXi AppTec, Pharmaron Ltd., Syngene International Ltd., Dalriada Drug Discovery, and Neurodex Inc. for their invaluable contributions to this work.

## Data availability

Supporting data and experimental methods are provided in the Supplementary Information and as Supplementary Tables.

## Notes

The authors declare the following competing financial interests. AJB, AJD, MM, BT, RM, BP, DK, JWK, PSH and EP are current employees of Dunad Therapeutics. LSC, ASAA, BWD, LRB, BL, EWS, SB, RT, GLS and LD are former employees of Dunad Therapeutics.

