## Supplementary Data and Methods for "Chemoproteomics discovery of a CNS-penetrant covalent inhibitor of PIKfyve"

### 1. Materials and Methods

#### Cell culture

HEK 293T cells (ATCC, CRL-3216) were cultured in DMEM supplemented with 10% v/v FBS. SU-DHL-6 cells (ATCC, CRL-2959) were cultured in RPMI-1640 medium supplemented with 10% v/v FBS. HCT-116 cells were cultured in McCoy's 5a medium supplemented with 10% v/v FBS. Cells were maintained in an incubator at 37 °C with 5% $CO_2$, passaging as appropriate according to the supplier's guidelines. iPSC-derived human motor neurons (BrainXcell, BX-0100 lot: 240923B, 240530H) were thawed and cultured as per supplier's protocol. In brief, cells were seeded on Poly-D-lysine-coated culture plates in base medium supplemented with 1x BrainFast Motor and pre-diluted Cultrex. On day 1 and 4, medium was replaced with base medium supplemented with BrainFast motor and BrainFast SK. On day 7, medium was replaced with base medium and cells were dosed as described in the indicated assay section.

#### Cysteine activity-based protein profiling (ABPP)

**TMT-based ABPP (library experiments):** Frozen cell pellets (approx. $5 \times 10^6$ cells) were thawed on ice and suspended in 500 µL of cold lysis buffer (50 mM 2-[4-(2-hydroxyethyl)piperazin-1-yl]ethane-1-sulfonic acid (HEPES) pH 7.5, 300 mM NaCl, 1.5 mM $MgCl_2$, 0.5% v/v NP-40, 5% v/v glycerol, 1 mM 3,3′,3″-phosphanetriyltripropanoic acid (TCEP), 1x cOmplete protease inhibitor (Roche)). Samples were lysed by probe sonication (15 seconds total, 30% amplitude, on ice) and clarified by centrifugation at 15,000 × *g* for 15 min at 4 °C. The supernatant was collected, the protein concentration was determined by BCA assay (Thermo Fisher Scientific), and samples were normalized to 500 µg protein input at 2 mg/mL concentration. Desthiobiotin-iodoacetamide (DTB-IA; 150 µM final concentration; MedChemExpress) was added to each sample and the alkylation reaction was left to proceed with mechanical agitation for 1 h at 25 °C, protected from light. The samples were sequentially treated with dithiothreitol (DTT; 5 mM final concentration for 30 min at 37 °C), and iodoacetamide (15 mM final concentration for 30 min at 25 °C, protected from light). Methanol/chloroform precipitation was performed, and the resultant protein pellet was then solubilized in 8 M urea (freshly prepared). 150 µg of protein input per sample was then diluted with 100 mM triethylammonium bicarbonate (TEAB; freshly prepared) to a final urea concentration of 0.5 M. Protein digestion was performed by addition of trypsin/LysC (1:40 trypsin/LysC:protein) and allowed to proceed overnight at 37 °C. Digested peptides were labeled using TMTpro16-plex reagents (Thermo Fisher Scientific), following the manufacturer's instructions. Following quenching with hydroxylamine (0.4% v/v, 15 min, RT), equal volumes of each sample were pooled and dried by SpeedVac (Thermo Fisher Scientific). Peptides were resuspended in PBS and added to pre-equilibrated high-capacity streptavidin agarose beads (90 µL slurry per pulldown; Thermo Fisher). The pulldown was left to proceed at 25 °C for 2 h with end-over-end rotation. The beads were sequentially washed with PBS, PBS + 0.1% w/v SDS, PBS and $H_2O$. Desthiobiotin-modified peptides were eluted by the sequential addition of $H_2O$ + 0.1% v/v trifluoroacetic acid (TFA), acetonitrile + 0.1% v/v TFA, and 50:50 v/v $H_2O$:acetonitrile + 0.1% v/v TFA. Eluates were combined and dried by SpeedVac. High-pH fractionation was then performed, sequentially eluting with 11, 14, 17, 19, 21.5, and 50% v/v acetonitrile in 0.1% v/v triethylamine, and dried by SpeedVac. Samples were reconstituted in 0.1% formic acid in $H_2O$ (solvent A) with sonication and injected onto an Aurora Ultimate C18 column (250 mm × 75 µm, 1.7 µm particle size). Peptides were separated across a 180 min gradient from 0-95% solvent B (80% acetonitrile, 0.1% formic acid in $H_2O$) at 50 °C and a flow rate of 0.3 µl/min, using a Vanquish Neo UHPLC in line with an Orbitrap Fusion Lumos mass spectrometer (Thermo Fisher Scientific). The Solvent B content across the gradient was 5% at 2 min, 11% at 19 min, 30% at 134 min, 48% at 164 min, and 95% at 165 min. MS1 spectra were collected in a 60,000 resolution MS1 scan (scan range 350-1,600 *m/z*, maximum injection time = 50 ms). MS2 spectra were acquired (precursor selection: mass range 400-1,500 *m/z*, intensity threshold = $1 \times 10^5$, dynamic exclusion of mono- and polyisotopic precursor ions within a mass tolerance of 10 ppm was used for 60 sec) with HCD fragmentation (collision energy = 35%, orbitrap resolution = 15,000, normalized AGC target = 100%, maximum injection time = 50 ms, isolation window = 1.1 *m/z*). Real-time search (RTS) was performed to select ions for synchronous precursor selection-MS3 quantification, requiring TMTpro as a fixed modification at Lys and peptide N-termini (304.2071 Da) and carbamidomethylation and the DTB-IA adduct as variable modifications at Cys (57.0215 and 296.1848 Da, respectively). The maximum search time was 40 ms. The MS3 scan was performed with HCD fragmentation (collision energy = 55%, orbitrap resolution = 50,000, scan range 100-500 *m/z*, maximum injection time = 100 msec). Data processing was performed in Proteome Discoverer (Thermo Fisher Scientific) with default parameters, including TMTpro as a fixed modification at Lys and peptide N-termini (304.2071 Da) and carbamidomethylation and the DTB-IA adduct as variable modifications at Cys (57.0215 and 296.1848 Da, respectively). The search was performed using the latest available canonical *Homo sapiens* UniProt FASTA database, obtained from Uniprot.org. Reporter ion intensities were corrected across TMT channels for isotopic impurities, as provided in the manufacturer's documentation for each reagent lot. Competition ratios (CR) were determined by taking an average of the intensities in the DMSO-treated control channels and dividing by the intensity in the compound-treated channels.[1] CR values were capped at a maximum of 20, interpreted as ≥ 95% target occupancy.

**DIA-based ABPP (DUN'058 Cys selectivity profiling):**

Cell pellets and sample processing were performed as above, with the following differences: (1) A biotin polyethyleneoxide iodoacetamide probe (BPI; Sigma Aldrich, B2059) was used at the same concentration in place of DTB; (2) No TMT labeling was performed, and samples were processed for individual injections across a dose range of DUN'058. Digested and desalted peptides were reconstituted in solvent A with sonication and injected onto an Aurora Ultimate C18 column (250 mm × 75 µm, 1.7 µm particle size). Peptides were separated across a 90 min gradient from 0 - 95% solvent C (90% acetonitrile, 0.1% formic acid in $H_2O$) at 50 °C and a flow rate of 0.3 µl/min, using a NanoElute LC system (Bruker). The Solvent C content across the gradient was 5% at 2 min, 22% at 30 min, 40% at 75 min, and 90% at 80 min. Samples were analyzed in DIA mode on a TimsTOF Pro 2 mass spectrometer (Bruker) in two separate injections. Injection 1 was performed with a 60,000 resolution MS1 scan (500-800 *m/z*), followed by MS/MS across 10 *m/z* windows at 30,000 resolution, with a normalized collision energy (NCE) setting of 28. Injection 2 was performed with a 60,000 resolution MS1 scan (800-1,150 *m/z*), followed by MS/MS across 10 *m/z* windows at 30,000 resolution, with an NCE setting of 28. Data were searched with DIA-NN using the latest available canonical *Homo sapiens* UniProt FASTA database, obtained from Uniprot.org, with default settings for library free search and adding carbamidomethylation and the BPI adduct as variable modifications at Cys. Data processing was performed by requiring the peptide to be quantified in all DMSO-treated samples (N = 4) and in both lowest concentration (0.064 nM) samples (N = 2). With this filtering performed, all remaining missing values were set to an arbitrary intensity value of 1 and CR values were calculated as above. Peptides that returned a $TO_{50}$ < 1 µM across the DUN'058 dose response were designated as hits, after fitting to a 4-parameter logistic model in GraphPad Prism 10.

**Kinase selectivity profiling**

Wild-type and lipid kinase selectivity analyses for DUN'465 and DUN'058 were performed at Reaction Biology Corporation (Malvern, PA) using the HotSpot™ Kinase Assay platform[2] and ADP-Glo (Promega), respectively. The HotSpot™ Kinase Assay was performed by preincubating test compounds (1 µM) at room temperature with the kinase, the substrate, and any appropriate cofactors for 20 minutes in the reaction buffer (20 mM HEPES pH 7.5, 10 mM $MgCl_2$, 1 mM EGTA, 0.01% Brij 35, 0.02 mg/mL BSA, 0.1 mM $Na_3VO_4$, 2 mM DTT, 1% DMSO). The reaction was initiated via the addition of 10 µM radioisotopically-labelled ATP ($^{32}$P-ɣ-ATP) and incubated for 2 h at room temperature. Kinase activity was detected using the P81 filter-binding method. Results were calculated as % remaining activity in test samples compared to DMSO controls.

Lipid kinase activity was measured using the ADP-Glo kinase assay kit (Promega #V9103). Test compounds (1 µM) were preincubated with each of the recombinant PI3K isoforms (α, β, δ, γ) and $PI(4,5)P_2$:PS substrate for 20 minutes. ATP (10 µM) was added to initiate the enzymatic reaction and the assay was incubated for 60 minutes at 30 °C, before addition of the ADP-Glo reagent (40 minutes incubation). Kinase detection reagent was added and incubated for 30 min and luminescence was detected using an Envision plate reader (Revvity). Results were calculated as % remaining activity in test samples compared to DMSO controls.

**Protein expression and purification**

**Human PIKfyve** (NP_055855.2; amino acids 1-2098) wild-type ($PIKfyve^{WT}$) and C1970A mutant ($PIKfyve^{C1970A}$) were separately cloned into a pcDNA3.1 mammalian expression vector with N-terminal 3xFLAG tag and C-terminal hexahistidine (His) tag. $PIKfyve^{WT}$ and $PIKfyve^{C1970A}$ constructs were expressed in Expi293F mammalian cells at 30 °C in Transpro CD 01 medium (Duoning). Cells were harvested after 72 h by centrifugation and flash-frozen for storage at -80 °C. Cell pellets were resuspended in cold PIKfyve lysis buffer (50 mM HEPES pH 8.0, 500 mM NaCl, 10% v/v glycerol, 0.1% Tween 20, 1 mM TCEP, 0.5% v/v protease inhibitor cocktail (Thermo Fisher Scientific), 0.1 mg benzonase/1 L culture) and lysed with 10 strokes of a Dounce homogenizer. The lysate was clarified by refrigerated centrifugation and the supernatant was incubated with anti-FLAG-G1 affinity resin (GenScript) for 1.5 h. The resin was washed three times with 10 bed volumes of PIKfyve wash buffer (50 mM HEPES pH 8.0, 10% v/v glycerol, 300 mM NaCl, 1 mM TCEP) and eluted three times with 2 bed volumes of PIKfyve elution buffer (PIKfyve wash buffer supplemented with 300 µg/mL FLAG peptide). Elution fractions were pooled and further purified using a HiLoad 16/600 Superose 6 pg (Cytiva) gel filtration column pre-equilibrated with PIKfyve storage buffer (50 mM HEPES pH 8.0, 600 mM NaCl, 5% v/v glycerol, 0.5 mM TCEP). Pure fractions were identified by SDS-PAGE, pooled, flash frozen in single-use aliquots, and stored at -80 °C. The presence of monomeric protein after one freeze-thaw cycle was confirmed by analytical gel filtration chromatography.

**Human Caspase 3** (Casp3; NP_004337.2; amino acids 1-277) was cloned into a pBAC-1 insect expression vector with an N-terminal cleavable maltose binding protein (MBP)-Sumo-3C tag and C-terminal His-HiBiT tag. Casp3 was expressed in baculovirus-infected Sf9 insect cells grown in HyClone™ SFX-Insect cell culture media (Cytiva). Cells were harvested by centrifugation 66 h after infection and flash frozen for storage at -80 °C. Frozen cell pellets were resuspended in ice-cold Casp3 lysis buffer (50 mM 2-amino-2-(hydroxymethyl)propane1,3-diol (Tris)-HCl pH 8.0, 300 mM NaCl, 5% v/v glycerol, 1 mM TCEP, 20 mM imidazole, 1% v/v protease inhibitor cocktail

(Thermo Fisher Scientific)) lysed with 10 strokes of a Dounce homogenizer. The lysate was clarified by refrigerated centrifugation and the supernatant was incubated with HisPur™ nickel-nitriloacetic acid (Ni-NTA) resin (Cytiva) for 0.5 h. After washing with 50 mL Casp3 wash buffer (50 mM Tris-HCl pH 8.0, 300 mM NaCl, 5% v/v glycerol, 1 mM TCEP, 20 mM imidazole), fractions were collected from a linear gradient of Casp3 wash buffer and Casp3 elution buffer (50 mM Tris-HCl pH 8.0, 300 mM NaCl, 5% v/v glycerol, 1 mM TCEP, 500 mM imidazole). The eluted sample was treated with 3C protease and dialyzed overnight against Casp3 wash buffer to remove the imidazole. The dialyzed sample was then passed through Ni-NTA resin to isolate the His-tagged Casp3 product. Pooled elution fractions were further purified by SEC on a Superdex S75 column (GE Healthcare) pre-equilibrated with Casp3 storage buffer (50 mM Tris-HCl pH 8.0, 300 mM NaCl, 1 mM TCEP). Peak fractions were pooled, analyzed by SDS-PAGE, flash-frozen in liquid nitrogen, and stored at -80 °C. Quality control after one freeze-thaw cycle showed purity of >90% that eluted as one main peak from an analytical SEC column, indicating no aggregation or oligomerization. Casp3 activity results from a self-cleavage event between Asp175 and Ser176, generating an N-terminal p17 subunit and a C-terminal p12 subunit.[3] Intact mass analysis confirmed the integrity of p12 subunits and partial degradation of p17 subunits (~39% showed excision of 6 C-terminal residues [CGIETD]). Peptide mapping confirmed the protein identity of both p17 and p12 subunits with high sequence coverage (94% and 97%, respectively).

**Human PIP4K2A** (NP_005019.2; amino acids 35-405) with 6 mutations (P176A, F178I, T196M, E229C, L230D, and I360Y; denoted as PIP4K2A$^{6mut}$) was cloned into a pET28a *E. coli* expression vector with an N-terminal His-TEV tag. PIP4K2A$^{6mut}$ was expressed in BL21-CodonPlus (DE3)-RIPL *E. coli* cells grown in TB medium supplemented with kanamycin (50 µg/mL) at 37 °C to an $OD_{600}$ of 0.8, after which protein expression was induced by addition of 0.25 mM IPTG. Cells were harvested by centrifugation after an additional 18 hours of growth (200 RPM, 18 °C). The cells were resuspended in PIP4K2A lysis buffer (20 mM $NaH_2PO_4$ pH 8.0, 300 mM NaCl, 10% v/v glycerol, 25 mM imidazole, 1% v/v protease inhibitor cocktail (Thermo Fisher Scientific), 0.1 mg benzonase/1 L culture) and lysed by high pressure homogenization (1000 bar, 3 passes). The lysate was clarified by refrigerated centrifugation and the supernatant was incubated with HisPur™ Ni-NTA resin for 1 h. After washing with 50 mL PIP4K2A wash buffer (20 mM Tris-HCl pH 8.0, 500 mM NaCl, 10% v/v glycerol, 2 mM TCEP, 25 mM imidazole), fractions were collected from a linear gradient of PIP4K2A wash buffer and PIP4K2A elution buffer (20 mM Tris-HCl pH 8.0, 500 mM NaCl, 10% v/v glycerol, 2 mM TCEP, 400 mM imidazole). Fractions of interest were pooled and further purified by SEC on a Superdex 200 column (GE Healthcare) in PIP4K2A low salt buffer (20 mM Tris-HCl pH 8.0, 250 mM NaCl, 2 mM TCEP). The protein was further purified by anion exchange chromatography using a HiTrap Q HP (Cytiva) and eluted with a gradient of PIP4K2A low salt buffer and PIP4K2A high salt buffer (20 mM Tris-HCl pH 8.0, 1 M NaCl, 2 mM TCEP). Fractions of interest were pooled, dialyzed overnight into PIP4K2A storage buffer (20 mM HEPES pH 8.0, 500 mM NaCl, 5% v/v glycerol, 2 mM TCEP), concentrated to > 10 mg/mL, flash-frozen in liquid nitrogen, and stored at -80 °C. Intact mass and peptide mapping analyses (92% sequence coverage) confirmed the identity of the protein.

**PIKfyve enzymatic inhibition assay (ADP-Glo)**

The inhibition of recombinant $PIKfyve^{WT}$ and $PIKfyve^{C1970A}$ was assessed using the ADP-Glo kinase assay kit (Promega #V9102). 300 nL of compounds (dissolved in DMSO) in serial dilution were transferred to 384-well Optiplates (Revvity #6007290) using an Echo 650 series liquid handler (Beckman Coulter). Recombinant $PIKfyve^{WT}$ or $PIKfyve^{C1970A}$ (5 µL) in the enzymatic assay buffer (25 mM HEPES pH 7.5, 10 mM $MgCl_2$, 1 µM $CaCl_2$, 2 mM DTT, 0.05% BSA, and 0.002% Triton-X100) was added to the compounds, sealed with TopSeal-A PLUS (Revvity #6050185) and incubated for 120 minutes at room temperature. 5 µL of ATP and PI(3)P:PS substrate was then added. The final concentrations of recombinant protein, ATP, and substrate were 5 nM, 20 µM, and 62.5 to 500 µM, respectively. The plate was resealed and incubated for 90 minutes at room temperature, then 10 µL of the ADP-Glo™ reagent was added and incubated for 60 minutes. Kinase detection reagent (20 µL) was added and incubated for 30 minutes, after which the luminescence (LUM) was detected using a Pherastar microplate reader (BMG Labtech). Controls included 0% inhibition (3% DMSO) and 100% inhibition (10 µM YM-201636; MedChemExpress #HY-13228). Percent inhibition was calculated as:

$$\%\text{Inhibition} = 100 \times \left[1 - \frac{\left(\text{LUM}_{\text{cmpd}} - \text{LUM}_{\text{pos}}\right)}{\left(\text{LUM}_{\text{neg}} - \text{LUM}_{\text{pos}}\right)}\right]$$

where $LUM_{cmpd}$, $LUM_{pos}$, and $LUM_{neg}$ represent the relative luminescence values of compound, YM-201636, and DMSO controls, respectively. $IC_{50}$ values were derived by fitting dose-response curves to a four-parameter logistic model:

$$Y = \text{Min} + \left[\frac{(\text{Max} - \text{Min})}{1 + (X/\text{IC}_{50})^{\text{Hill slope}}}\right]$$

or a three-parameter model if the Max or Min of the curves were fixed to 100% or 0%, respectively. $IC_{50}$ curves were plotted using GraphPad Prism 10.

The ATP $K_M^{app}$ for PIKfyve$^{WT}$ and PIKfyve$^{C1970A}$ were determined by titrating ATP (1.95 - 125 μM) into a reaction mixture containing 5 nM of enzyme, 62.5 to 500 μM of PI(3)P:PS, and 3% DMSO in the enzymatic assay buffer. 5 μL of enzyme was preincubated with DMSO for 2 hours before initiating the reaction with 5 μL of ATP and PI(3)P:PS. After 90 minutes, the luminescence values were determined using the ADP-Glo kinase assay kit (Promega #V9102). ATP $K_M^{app}$ was determined by fitting the data to a "One site – specific binding model", [ $\mathrm{RLU} = \frac{RLU_{max}\times[ATP]}{[ATP]+K_M}$ ], using GraphPad Prism 10.

#### Selective proteolysis mediated intact mass analysis (IMA)

Recombinant Casp3 (20.5 nM final concentration) was added to a solution of recombinant PIKfyve$^{WT}$ or PIKfyve$^{C1970A}$ protein (410 nM final concentration) in PIKfyve storage buffer. Casp3 digestion was performed overnight at 25 °C. The digested PIKfyve protein was dispensed directly into 96-well or 384-well plates containing a 20x stock of the compound of interest as a DMSO solution. The final %/v DMSO in the assay was 5%. The reaction was allowed to proceed for the indicated time at 25 °C, then quenched by the addition of 0.5% v/v formic acid (final concentration). The samples were diluted 3-fold with LC-MS grade $H_2O$ and equilibrated to 4 °C before injection onto a RapidFire system in-line with an ESI-ToF mass spectrometer (Agilent). Raw spectra were analyzed using MassHunter BioConfirm software (Agilent). The % adducts were reported by quantifying the deconvoluted signal intensity of the unmodified protein (35,913 Da) and any observed covalent adducts (single, double modification etc).

#### Peptide mapping

Recombinant PIKfyve$^{WT}$ protein (410 nM) was treated with DUN'465 or DUN'058 (2.05 μM) in PIKfyve storage buffer at 25 °C for 1 h. The sample was then processed sequentially through acetone precipitation, reduction (dithiothreitol), and alkylation (iodoacetamide) before overnight digestion with trypsin/LysC (1:40 trypsin/LysC:protein) at 37 °C. Digested peptides were purified by stage tip purification, resuspended in $H_2O$ + 0.1% v/v formic acid, and injected onto a Thermo QE-HF instrument. Data analysis was performed using PEAKS software, including appropriate covalent adducts for DUN'465 (parent molecular formula = $C_{26}H_{27}ClF_5N_7O_2S$; formal adduct molecular formula = $C_{26}H_{26}ClF_4N_7O_2S$) and DUN'058 (parent molecular formula = $C_{27}H_{31}FN_6O_2$; formal adduct molecular formula = $C_{27}H_{31}FN_6O_2$) as variable modifications at cysteine.

**Determination of $k_{inact}$ and $K_I$ by IMA**

Samples were prepared by selective proteolysis IMA as described above. 100 nM PIKfyve$^{WT}$ was treated with 30, 100, 300, and 900 nM DUN'058 for 10, 30, 60, 90, 180, 270, and 360 min at 25 °C. Samples were injected into a Waters QToF mass spectrometer operating in positive ion mode with a scan range of 300-3,500 *m/z.* A 15-min gradient of 0-100% solvent B (90% acetonitrile in $H_2O$ with 0.1% FA) was employed. The % adducts were reported by quantifying the deconvoluted signal intensity of the unmodified protein (35,913 Da) and singly adducted protein (36,404 Da). Occupancy of PIKfyve C1970 by DUN'058 was plotted as a function of time and the observed rate constant ($k_{obs}$) at each concentration was determined by fitting the curves using mono-exponential association and constraining occupancy to zero at time zero. $k_{obs}$ values were then plotted as a function of DUN'058 concentration, and $k_{inact}$ and $K_I$ values were determined by fitting to the equation $k_{obs} = \frac{k_{inact} \times [DUN'058]}{[DUN'058] + K_I}$.

**PIP4K2A$^{6mut}$-DUN'058 crystallization**

Recombinant PIP4K2A$^{6mut}$ protein (10 mg/mL in PIP4K2a storage buffer) was mixed with DUN'058 (50 mM in 100% DMSO) to reach a final compound concentration of 1 mM. The mixture was incubated for 6 h at room temperature before desalting treatment using Zeba Spin desalting columns (Thermo Fisher Scientific). Confirmation of 100% single adduct modification was determined by IMA. Crystals of PIP4K2A$^{6mut}$ with DUN'058 were obtained by co-crystallization at 4 °C with the sitting-drop vapour diffusion method using a mixture of 0.1 µL adduct-protein solution, 0.02 µL seed solution and 0.18 µL reservoir solution containing 15% PEG 3350, 200 mM potassium formate, and 100 mM HEPES pH 8.0. Plate-shaped crystals typically appeared within 2 days and were harvested after 7 days. Crystals were cryo-protected by plunging into reservoir solution supplemented with 20% v/v glycerol prior to flash-freezing in liquid nitrogen. Crystals were stored in liquid nitrogen until data collection.

**PIP4K2A$^{6mut}$-DUN'058 co-crystal structure determination**

Datasets of PIP4K2A$^{6mut}$ co-crystals were collected on a PILATUS3 6M detector at the Shanghai Synchrotron Radiation Facility (31124.02.SSRF) on the BL19U1 Protein Complex Crystallography beamline using a wavelength of 0.9786 Å with crystals maintained at approximately 100 K. The data were processed using autoPROC[4] and indexed into space group $P4_22_12$. Diffraction was anisotropic and was processed with STARANISO. The anisotropic diffraction limits of the unit cell were a = b = 3.16 Å, c = 4.06 Å. Structure determination was performed by molecular replacement in Phaser[5] using a previously published PIP4K2A$^{WT}$ kinase domain structure (PDB: 2ybx) as a search model. After manual ligand placement, iterative cycles of manual model building and automated refinement were performed using COOT[6] and Refmac5[7], respectively. The Ramachandran statistics for the final structure were: 91.7% favoured, 8.3% allowed, and 0% outliers. The co-crystal structure of PIP4K2A$^{6mut}$ bound to DUN'058 has been deposited into the Protein Data Bank with PDB ID 9zmo. Data collection and refinement statistics are shown in Supplementary Table 3.

**Target occupancy by PRM-based targeted mass spectrometry**

**Cell pellet processing:** Frozen cell pellets (approx. 1-2 × $10^6$ cells) were thawed on ice and suspended in 200 µL of cold lysis buffer (phosphate buffered saline (PBS) pH 7.4, 0.1% v/v NP-40, 1x cOmplete protease inhibitor). Samples were lysed by probe sonication (60 seconds total at 30% amplitude, on ice) and clarified by centrifugation at 12,000 × *g* for 10 min at 4 °C. The supernatant was collected and the protein concentration was determined by BCA assay (Thermo Fisher Scientific). 100 µg protein input was treated sequentially with DTT and iodoacetamide, before processing and digestion with S-Trap spin columns following the manufacturer's instructions.

**Brain homogenate spike-in experiments:** Cold lysis buffer (50 mM HEPES pH 7.5, 150 mM NaCl, 0.05% v/v Triton X-100, 10% v/v glycerol, 1 mM TCEP, 1x cOmplete protease inhibitor (Roche)) was added to mouse brain tissue (20 mg with 1 mL lysis buffer, or right brain with 3 mL lysis buffer). Tissue was homogenized by mechanical bead homogenization using a TissueLyser II instrument, followed by probe sonication (60 seconds total at 30% amplitude, on ice). Samples were clarified by centrifugation at 12,000 × *g* for 10 min at 4 °C. The supernatant was collected and protein concentration was determined by BCA assay. Aliquots of 100 µg brain homogenate were treated with a dose response of DUN'058 for 1 h at 25 °C. Excess compound was then removed by acetone precipitation. 20 µg protein input was treated sequentially with DTT and iodoacetamide, before processing and digestion with S-Trap spin columns following the manufacturer's instructions.

***In vivo* tissue samples:** One mL of cold lysis buffer (1x RIPA buffer, 1x cOmplete protease inhibitor (Roche)) was added to 20 mg of mouse brain tissue. Tissue was homogenized by mechanical bead homogenization using a TissueLyser II instrument, followed by probe sonication (60 seconds total at 30% amplitude, on ice). Samples were clarified by centrifugation at 12,000 × *g* for 10 min at 4 °C. The supernatant was collected and the protein concentration was determined by BCA assay. 100 µg protein input was treated sequentially with DTT and iodoacetamide, before processing and digestion with S-Trap spin columns following the manufacturer's instructions.

**PRM-based targeted mass spectrometry:** Peptides were reconstituted in solvent A with sonication and injected onto an Aurora Ultimate C18 column (250 mm × 75 µm, 1.7 µm particle size) in line with an Orbitrap Eclipse Tribrid or Q Exactive HF Orbitrap mass spectrometer. Peptides were separated across a 60 min gradient from 0-90% solvent C (90% acetonitrile, 0.1% formic acid in $H_2O$) at 50 °C and a flow rate of 0.35 µL/min. The Solvent C content across the gradient was 5% at 5 min, 25% at 45 min, 40% at 55 min, and 90% at 60 min. Parallel reaction monitoring-based targeted MS was used to determine covalent target occupancy (TO) by monitoring the tryptic peptide containing PIKfyve C1970 (`ESC(+`carbamidomethylation`)DVVLLDENLLK`; 823.9189 *m/z*; +2 precursor charge; RT = 52 min). An additional PIKfyve peptide (`FTC(+`carbamidomethylation`)IDPIVLQER`; 745.8872 *m/z*; +2 precursor charge; RT = 45 min) was monitored to assess if the overall abundance of PIKfyve was affected by compound treatment. Four peptides from four housekeeping proteins (beta-tubulin (`ISVYYNEATGGK`; 651.3223 *m/z*; +2 precursor charge; RT = 26 min), alpha-actinin-4 (`LVSIGAEEIVDGNAK`; 757.9067 *m/z*; +2 precursor charge; RT = 39 min), alpha-actinin-1 (`VGWEQLLTTIAR`; 693.8907 *m/z*; +2 precursor charge ; RT = 56 min), GAPDH (`VGVNGFGR`; 403.2194 *m/z*; +2 precursor charge; RT = 22 min)) were quantified as standards for normalization. All peptides are conserved between human and mouse PIKfyve protein. The peptide containing the covalent adduct of DUN'058 with C1970 was added to the method to confirm direct covalency with PIKfyve C1970 (`ESC(+`DUN'058`)DVVLLDENLLK`; 694.0243 *m/z*; +3 precursor charge; RT = 57 min). C1970 TO% was determined using the following equation: $TO\%_{C1970} = 100 – (C1970\ AU_t / C1970\ AU_0)$, where C1970 $AU_t$ is the area signal from the C1970-containing peptide at time *t*, and $AU_0$ is the area signal from the C1970-containing peptide at time = 0 or from vehicle treated animals, determined by Xcalibur software (Thermo Fisher Scientific).

**NanoBRET target engagement assay**

HEK 293T cells were transiently transfected with a plasmid encoding PIKfyve-NanoLuc (Promega), plated into a 384-well assay plate, and incubated at 37°C for 24 h. Transfected cells were treated with NanoBRET TE intracellular kinase (K-8) probe (130 nM final concentration; Promega) and compound in a total volume of 30 µl Opti-MEM for 1 h at 37 °C. 15 µL of 3 × complete substrate plus inhibitor solution (Promega) was added and the cells were incubated for 2 min at RT. Luminescence was measured at the donor (450 nM) and acceptor (610 nM) emission wavelengths, using an EnVision plate reader (PerkinElmer). The BRET ratio was determined by dividing the acceptor emission value by the donor emission value and multiplying by 1,000. BRET ratios were normalized to high (1 µM apilimod) and low (DMSO) controls, and $IC_{50}$ values were derived by fitting to a 4-parameter logistic model. $IC_{50}$ curves were plotted using GraphPad Prism 10.

**Global proteomics**

Frozen cell pellets (approx. $2 \times 10^{6}$ cells) were thawed on ice and suspended in 200 µL of cold lysis buffer (1x RIPA buffer, 1x cOmplete protease inhibitor (Roche)). Samples were lysed by probe sonication (60 seconds total at 30% amplitude, on ice) and clarified by centrifugation at 12,000 × *g* for 10 min at 4 °C. The supernatant was collected, and the protein concentration was determined by BCA assay. 100 µg protein input was treated sequentially with DTT and iodoacetamide, before processing and digestion with S-Trap spin columns following the manufacturer's instructions. Peptides were reconstituted in solvent A with sonication and injected onto an Aurora Ultimate C18 column (250 mm × 75 µm, 1.7 µm particle size; IonOpticks). Peptides were separated across a 55 min gradient from 0-90% solvent C (90% acetonitrile, 0.1% formic acid in $H_2O$) at 50 °C and a flow rate of 0.3 µL/min, using a NanoElute LC system (Bruker). The Solvent C content across the gradient was 32% at 39 min, 40% at 43 min, 60% at 46 min, and 90% at 60 min. Samples were analyzed in DIA mode on a TimsTOF Pro 2 mass spectrometer (Bruker), acquiring a 60,000 resolution MS1 scan (100-1,700 *m/z*), followed by MS/MS across 40 windows at 30,000 resolution, with an NCE setting of 28. Data was searched with DIA-NN using the latest available canonical *Homo sapiens* UniProt FASTA database, obtained from Uniprot.org, with default parameters for library free search. Resulting protein-level intensities were processed to generate $Log_2$(Fold Change(DUN'058/DMSO)) values and *p*-values, which were adjusted for multiple hypothesis testing using the Benjamini-Hochberg method.[8] Proteins returning a $Log_2$(FC) of ≤ -0.5 or ≥ 0.5 and an adjusted *p*-value of < 0.05 were considered significant.

**TFEB phosphorylation assay**

SU-DHL-6 cells (45,000 cells per well) were seeded in 384 well plates in Opti-MEM + 1% v/v FBS. Compounds were added as DMSO solutions in a 7-point, 3.5-fold dilution series from 10 µM to 5.44 nM (final concentration) in triplicate. The plate was incubated at 37 °C for 2 h. Cells were lysed with the addition of 10 µL of 5x AlphaLISA lysis buffer (Revvity) and PI (Thermo Fisher Scientific) per well with shaking for 20 min at RT. In the absence of light, 10 µL of donor and acceptor bead mix was added to each well, prepared following the manufacturer's instructions, and the resulting solutions incubated overnight at 4 °C with shaking. The plate was then equilibrated to RT and analyzed on an EnVision™ plate reader (PerkinElmer) using default AlphaLISA parameters.

**Quantification of GPNMB mRNA by qPCR**

iPSC-derived hMN were plated (40,000 per well) in a 96-well plate, cultured for 7 days as described above, and then treated with the indicated concentrations of DUN’058 for 24 h in culture medium. cDNA was prepared with TaqMan Cells-to-CT Kit (Invitrogen, A35374), following the manufacturer’s instructions. qPCR reactions were performed using QuantStudio™ 7 (Thermo Fisher Scientific) with PrimeTime® Gene Expression Master Mix (IDT, 1055772) using primer/probes for GPNMB (ThermoFisher Scientific, Hs01095669_m1).

**PK studies**

**50 mg/kg oral PK study:** DUN’058 was dosed at 50 mg/kg by oral gavage to fasted male C57BL/6 mice (4 cohorts, n = 3) as a solution in DMSO:PEG400:TPGS:20% w/v aqueous HP-β-CD (10:20:10:60 v/v) at a concentration of 5 mg/mL. Blood was sampled via the saphenous vein at 0.5, 1, 2 and 4 h and brains were collected at 0.5, 1, 2 and 4 h. Plasma samples (5 µL) were precipitated with 180 µL of acetonitrile containing Verapamil as internal standard. Samples were vortexed at 1,000 rpm for 10 min and centrifuged for 10 min at 4,000 rpm. 120 µL of supernatant was transferred to a 96-well plate and analyzed using LC-MS/MS and compared against a standard curve with the compound. Homogenized brain samples (10 µL) were precipitated with 300 µL of acetonitrile containing Verapamil as internal standard. Samples were vortexed at 1,000 rpm for 10 min and centrifuged for 10 min at 4,000 rpm. 150 µL of supernatant was transferred to a 96-well plate and analyzed using LC-MS/MS and compared against a standard curve with the compound. Pharmacokinetic parameters were calculated using the non-compartmental analysis module of Phoenix WinNonlin (version 8.2).

**150 mg/kg oral PK/TO study:** DUN’058 was dosed at 150 mg/kg by oral gavage to fasted male C57BL/6 mice (5 cohorts, n = 3) as a suspension in DMSO:PEG400:TPGS:20% w/v aqueous HP-β-CD (10:20:10:60 v/v) at a concentration of 15 mg/mL. Blood was sampled at 0.25, 0.5, 2 and 4 h from one cohort of animals only, and brain, spleen, duodenum and colon were collected at 4, 24, 48, 96 and 120 h. Blood was collected into 1.5 mL Eppendorf tubes with EDTA-2K, then centrifuged at 2,500 × *g* at 4 °C for 15 min to obtain plasma for bioanalysis. Each brain and spleen were divided into 2 parts and preserved by snap freezing in liquid nitrogen, respectively. Brain and spleen homogenate was prepared by homogenizing brain and spleen tissue with 9 volumes w:v of PBS:MeOH (v:v = 2:1) buffer. Plasma, brain and spleen homogenate was quenched with 5 ng/mL Verapamil solution (100 µL) as internal standard and then vortex-mixed for 5 min at 800 rpm and centrifuged for 15 min at 3,220 × *g* (4,000 rpm) at 4 °C. The supernatant (30 µL) was diluted with $H_2O$ (30 µL) and fully vortexed for 5 min. The diluted solutions were directly injected for LC-MS/MS analysis and compared against a standard curve of 0.200-400 ng/mL of the compound in plasma, brain and spleen.

**5-Day mouse tolerability study**

DUN'058 was dosed once daily over 5 days at 50 mg/kg by oral gavage to fed male C57BL/6 mice (n = 6) as a solution in DMSO:PEG400:TPGS:20% w/v aqueous HP-β-CD (10:20:10:60 v/v) at a concentration of 5 mg/mL. Post last dose, blood was sampled at 0.5, 8 and 24 h, n = 3 for each timepoint, and brain and spleen were collected at 0.5 and 24 h, n = 3 for each timepoint. Blood was collected into 1.5 mL Eppendorf tubes with EDTA-2K, then centrifuged at 2,500 × *g* at 4 °C for 15 min to obtain plasma for bioanalysis. Each brain and spleen were divided into 2 parts and preserved by snap freezing in liquid nitrogen, respectively. Brain and spleen homogenate was prepared by homogenizing brain and spleen with 9 volumes w:v of PBS:MeOH (v:v = 2:1) buffer. Plasma, brain and spleen homogenate was quenched with 5 ng/mL Verapamil solution (100 µL) as internal standard and then vortex-mixed for 5 min at 800 rpm and centrifuged for 15 min at 3,220 × *g* (4,000 rpm) at 4 °C. The supernatant (30 µL) was diluted with $H_2O$ (30 µL) and fully vortexed for 5 min. The diluted solutions were directly injected for LC-MS/MS analysis and compared against a standard curve of 1-1,000 ng/mL of the compound in plasma for the 0.5 h plasma samples, and 0.1-100 ng/mL of the compound in plasma, brain and spleen for the 8 h and 24 h plasma, brain and spleen samples.

**Compound Synthesis**

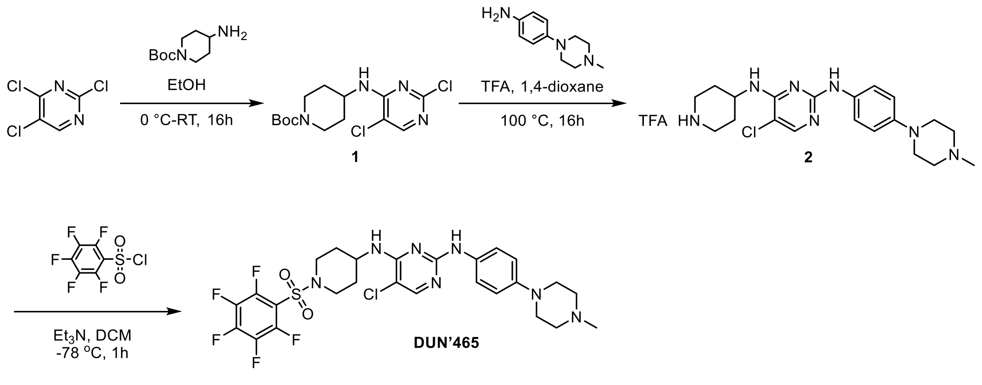

***tert*-butyl 4-((2,5-dichloropyrimidin-4-yl)amino)piperidine-1-carboxylate (1):**

To an oven-dried 500 mL two-neck round-bottom flask equipped with $N_2$ balloon and a rubber septum 2,4,5-trichloropyrimidine (10.0 g, 54.5 mmol) and ethanol (100 mL) were added. The resulting solution was cooled to 0 °C and *tert*-butyl 4-aminopiperidine-1-carboxylate (16.38 g, 82 mmol) was added. The reaction mixture was stirred under a nitrogen atmosphere for 16 h at 50 °C. The progress of the reaction was monitored by LC-MS. After completion of the reaction, ice water (100 mL) was added and the mixture was extracted with ethyl acetate (2 × 80 mL). The combined organic portions were washed with water (2 × 50 mL) and a saturated sodium chloride solution (50 mL). The organic portion was dried over anhydrous $Na_2SO_4$, filtered, and concentrated under reduced pressure. The crude residue was purified by silica gel flash column chromatography using 30-40 % ethyl acetate in hexanes to provide *tert*-butyl 4-((2,5-dichloropyrimidin-4-yl)amino)piperidine-1-carboxylate (8.10 g, 22.63 mmol, 42 %) as an off-white solid. **MS (ES+):** *m/z* = 347/349 $[M+H]^+$.

**5-chloro-*N*$^2$-(4-(4-methylpiperazin-1-yl)phenyl)-*N*$^4$-(piperidin-4-yl)pyrimidine-2,4-diamine trifluoroacetate salt (2):**

To an oven dried 25 mL two-neck round-bottom flask equipped with $N_2$ balloon, TFA (2.399 mL, 31.1 mmol) was added to a solution of *tert*-butyl 4-((2,5-dichloropyrimidin-4-yl)amino)piperidine-1-carboxylate (**1**, 0.2 g, 0.576 mmol) and 4-(4-methylpiperazin-1-yl)aniline (0.132 g, 0.691 mmol) in 1,4-dioxane (2 mL). The resulting reaction mixture was stirred at 100 °C for 16 h. After the reaction was deemed complete by TLC, the volatiles were removed under reduced pressure to afford 5-chloro-*N*$^2$-(4-(4-methylpiperazin-1-yl)phenyl)-*N*$^4$-(piperidin-4-yl)pyrimidine-2,4-diamine trifluoroacetate salt (0.260 g, 0.647 mmol, 90 %). **MS (ES+):** *m/z* = 402/404 $[M+H]^+$.

**5-chloro-*N*$^2$-(4-(4-methylpiperazin-1-yl)phenyl)-*N*$^4$-(1-((perfluorophenyl)sulfonyl)piperidin-4-yl)pyrimidine-2,4-diamine (DUN'465):**

To an oven dried 50 mL two-neck round-bottom flask equipped with $N_2$ balloon, 5-chloro-*N*$^2$-(4-(4-methylpiperazin-1-yl)phenyl)-*N*$^4$-(piperidin-4-yl)pyrimidine-2,4-diamine trifluoroacetate salt (**2,** 0.250 g, 0.622 mmol), triethylamine (0.874 mL, 6.22 mmol) and dichloromethane (3 mL) were added and the mixture was stirred for 10 min at 0 °C. The reaction mixture was cooled to -78 °C. A solution of 2,3,4,5,6-pentafluorobenzenesulfonyl chloride (0.166 g, 0.622 mmol) in dichloromethane (3 mL) was added and mixture stirred for 1 h at -78 °C. The reaction was quenched with ice water (3 mL) and extracted with dichloromethane (2 × 10 mL). The combined organic portions were dried over anhydrous sodium sulphate, filtered and concentrated under reduced pressure afforded the crude product. The crude product was purified by prep HPLC to afford 5-chloro-$N^2$-(4-(4-methylpiperazin-1-yl)phenyl)-$N^4$-(1-((perfluorophenyl)sulfonyl)piperidin-4-yl)pyrimidine-2,4-diamine (59 mg, 0.091 mmol, 15 %) as an off-white solid. **$^1$H-NMR (400 MHz, DMSO-*d₆*):** δ 8.97 (s, 1H), 7.90 (s, 1H), 7.51 (d, J = 9.2 Hz, 2H), 6.88 (d, J = 7.6 Hz, 1H), 6.81 (d, J = 8.8 Hz, 2H), 3.98-3.87 (m, 1H), 3.80 (d, J = 11.6 Hz, 2H), 3.06-2.99 (m, 4H), 2.87 (t, J = 11.8 Hz, 2H), 2.49-2.43 (m, 4H), 2.24 (s, 3H), 2.06 (d, J = 10.8 Hz, 2H), 1.78-1.65 (m, 2H). **$^{19}$F-NMR (376 MHz, DMSO-$d_6$):** δ -135.86 to -135.92 (2F), -147.16 to -147.28 (1F), -159.65 to -159.79 (2F). **MS (ES+):** *m/z* = 632/634 [M+H]$^+$.

**Preparative HPLC column and solvents:** Column: Shimadzu SHIMPACK GIST C18 (150 × 20.0 mm) 5 µm; mobile phase A: 10 mM $NH_4HCO_3$ in water; mobile Phase B: MeCN; flow rate: 15.0 mL/min.

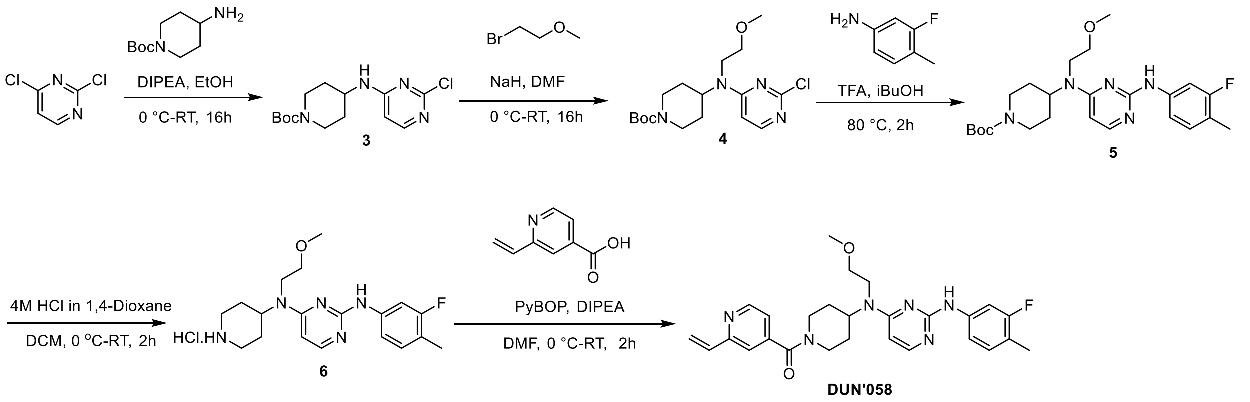

***tert*-butyl 4-((2-chloropyrimidin-4-yl)amino)piperidine-1-carboxylate (3):**

In an oven-dried 500 mL two-neck round-bottom flask equipped with $N_2$ balloon and a rubber cap, 2,4-dichloropyrimidine (15 g, 101 mmol) and EtOH (200 mL) were added. The reaction mixture was cooled to 0 °C, tert-butyl 4-aminopiperidine-1-carboxylate (20.17 g, 101 mmol) and DIPEA (13.01 g, 101 mmol) were introduced at the same temperature then the resulting reaction mixture was stirred for 16 h at 25 °C. The progress of the reaction was monitored by TLC. After completion of the reaction, the volatiles were removed under reduced pressure. The obtained crude mass was purified by flash column chromatography using 40 % EtOAc in hexane to afford *tert*-butyl 4-((2-chloropyrimidin-4-yl)amino)piperidine-1-carboxylate (20 g, 62.0 mmol, 62 %) as an off-white solid. **MS (ES+):** m/z = 313/315 [M+H]$^+$.

***tert*-butyl 4-((2-chloropyrimidin-4-yl)(2-methoxyethyl)amino)piperidine-1-carboxylate (4):**

To an oven dried 250 mL two-neck round-bottom flask equipped with $N_2$ balloon and a rubber cap, *tert*-butyl 4-((2-chloropyrimidin-4-yl)amino)piperidine-1-carboxylate (**3**, 6 g, 19.18 mmol) and DMF (100 mL) were added. The reaction mixture was cooled to 0 °C, sodium hydride (2.30 g, 57.5 mmol) was added portion wise at the same temperature, stirred for 20 min, then a solution of 1-bromo-2-methoxyethane (9.01 mL, 96 mmol) in DMF (10 mL) was introduced. The resulting reaction mixture was stirred for 16 h at 25 °C. The progress of the reaction was monitored by TLC and upon completion the reaction was quenched with ice water (100 mL) and extracted with ethyl acetate (2 × 100 mL). The combined organic layers were dried over anhydrous sodium sulphate, filtered and concentrated under reduced pressure to afford crude product. The obtained crude mass was purified by flash column chromatography using 50-60 % ethyl acetate in hexanes to afford tert-butyl 4-((2-chloropyrimidin-4-yl)(2-methoxyethyl)amino)piperidine-1-carboxylate (6.0 g, 16.21 mmol, 85 %) as an off-white solid.
**MS (ES+):** m/z = 371/373 $[M+H]^+$.

***tert*-butyl 4-((2-((3-fluoro-4-methylphenyl)amino)pyrimidin-4-yl)(2-methoxyethyl)amino)piperidine-1-carboxylate (5):**
To an oven dried 250 mL two-neck round-bottom flask equipped with $N_2$ balloon, *tert*-butyl 4-((2-chloropyrimidin-4-yl)(2-methoxyethyl)amino)piperidine-1-carboxylate (**4**, 4 g, 10.79 mmol), isobutanol (50 mL), 3-fluoro-4-methylaniline (1.35 g, 10.79 mmol) and TFA (1.23 g, 10.79 mmol) were added at room temperature. Then the reaction mixture was stirred for 2 h at 80 °C. The progress of the reaction was monitored by TLC. After completion, the reaction mixture was concentrated under reduced pressure to afford *tert*-butyl 4-((2-((3-fluoro-4-methylphenyl)amino)pyrimidin-4-yl)(2-methoxyethyl)amino)piperidine-1-carboxylate (4.2 g, 8.89 mmol, 82 %). The compound was used in the next step without additional purification. **MS (ES+):** *m/z* = 460 $[M+H]^+$.

***N*²-(3-fluoro-4-methylphenyl)-*N*⁴-(2-methoxyethyl)-*N*⁴-(piperidin-4-yl)pyrimidine-2,4-diamine hydrochloride (6):**
In an oven dried 50 mL round-bottom flask equipped with $N_2$ balloon, *tert*-butyl 4-((2-((3-fluoro-4-methylphenyl)amino)pyrimidin-4-yl)(2-methoxyethyl)amino)piperidine-1-carboxylate (**5**, 4 g, 8.47 mmol) and dichloromethane (30 mL) were combined, cooled in an ice bath and 4M HCl solution in dioxane (42.3 mL, 169 mmol) was added at 0 °C. The reaction mixture was stirred at 25 °C for 2 h. The progress of the reaction was monitored by TLC. After completion, the volatiles were removed under reduced pressure to afford a crude residue that was triturated with MTBE to give

*N*[2]-(3-fluoro-4-methylphenyl)-*N*[4]-(2-methoxyethyl)-*N*[4]-(piperidin-4-yl)pyrimidine-2,4-diamine hydrochloride (3.2 g, 8.05 mmol, 95 %) as an off-white solid. **MS (ES+):** *m/z* = 360 $[M+H]^+$.

**(4-((2-((3-fluoro-4-methylphenyl)amino)pyrimidin-4-yl)(2-methoxyethyl)amino)piperidin-1-yl)(2-vinylpyridin-4-yl)methanone (DUN'058):**

To an oven dried 50 mL two-neck round-bottom flask equipped with $N_2$ balloon, *N*[2]-(3-fluoro-4-methylphenyl)-*N*[4]-(2-methoxyethyl)-*N*[4]-(piperidin-4-yl)pyrimidine-2,4-diamine hydrochloride (2.5 g, 6.29 mmol), 2-vinylisonicotinic acid (1.39 g, 6.92 mmol), DIPEA (3.39 mL, 18.9 mmol) and DMF (30 mL) were added. The reaction mixture was cooled to 0 °C, PyBOP (4.91 g, 9.44 mmol) was introduced, and the mixture was stirred for 2 h at room temperature. The progress of the reaction was monitored by TLC and LC-MS. After the starting material was consumed, the reaction was quenched with ice water (25 mL) and extracted with ethyl acetate (2 × 50 mL). The combined organic layers were dried over anhydrous sodium sulphate, filtered, and concentrated under reduced pressure affording the crude product. The obtained crude was purified by preparative HPLC to afford (4-((2-((3-fluoro-4-methylphenyl)amino)pyrimidin-4-yl)(2-methoxyethyl)amino)piperidin-1-yl)(2-vinylpyridin-4-yl)methanone (1.2 g, 2.39 mmol, 38 %) as an off-white solid. **$^1$H-NMR (400 MHz, DMSO-*d*$_6$, 80 °C ):** δ 8.89 (s, 1H), 8.64 (d, J = 4.8 Hz, 1H), 7.97 (d, J = 6.0 Hz, 1H), 7.70-7.64 (m, 1H), 7.53 (s, 1H), 7.36-7.29 (m, 2H), 7.11 (t, J = 8.8 Hz, 1H), 6.92-6.83 (m, 1H), 6.32-6.26 (m, 1H), 6.22 (d, J = 6.0 Hz, 1H), 5.56-5.52 (m, 1H), 4.55 (br.s, 2H), 3.63-3.47 (m, 5H), 3.31 (s, 3H), 2.17 (s, 3H), 1.87-1.67 (m, 4H). Two protons obscured by water peak. **$^{19}$F-NMR (376 MHz, DMSO-*d*$_6$):** δ -116.96 (1F). **MS (ES+):** *m/z* = 491 $[M+H]^+$.

**Preparative HPLC column and solvents:** Column: Shimadzu SHIMPACK GIST C18 (250 × 20.0 mm) 5 µm; mobile phase A: 10 mM $NH_4HCO_3$ in water; mobile Phase B: MeCN; flow rate: 15.0 mL/min.

### 2. Supplementary Figures

**Supplementary Figure 1.** Cysteine profiling ABPP data for DUN'465. The experiment was run in HEK293T cells, treated with 10 µM DUN'465 for 2 h. Cutoffs displayed = $Log_2$ fold change ≤ 1 or ≥ 1, *p*-value < 0.01. Data is provided as Supplementary Table 2.

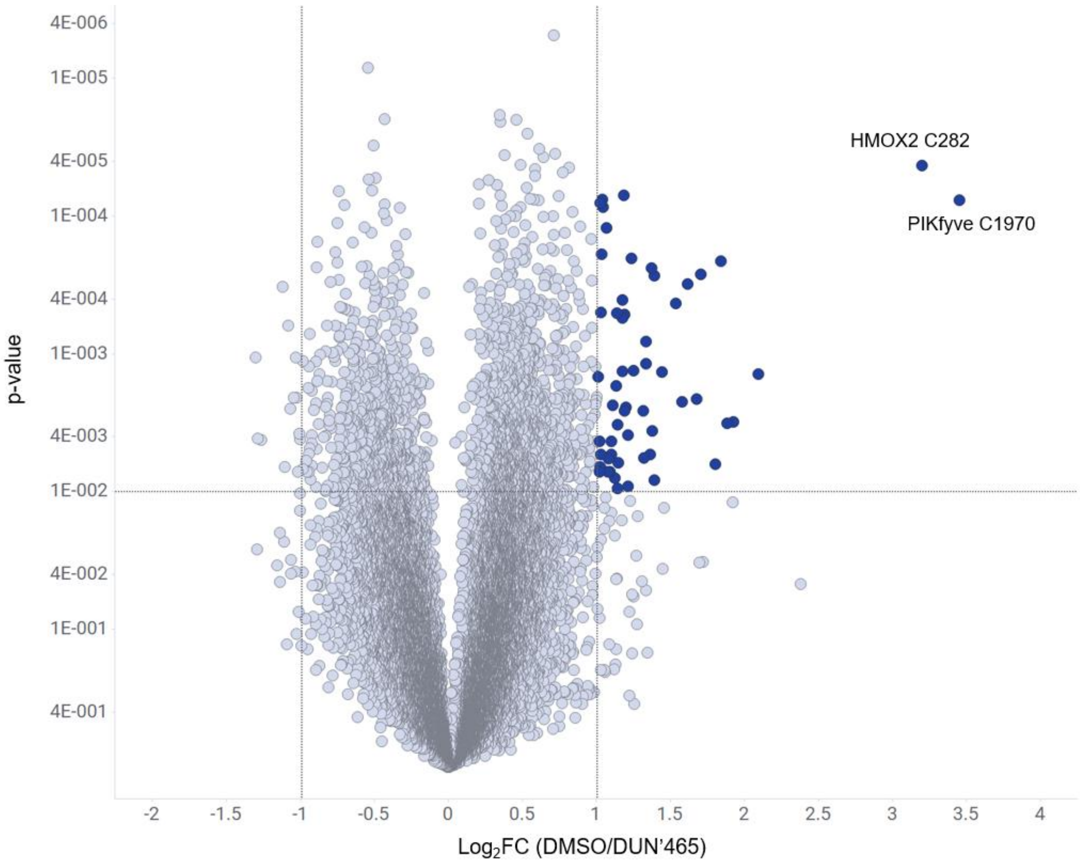

**Supplementary Figure 2.** LC-MS/MS analysis of trypsin-digested PIKfyve$^{WT}$ after treatment with DUN’465 (1 h, 25 °C) confirms covalent modification at C1970 (DUN’465 MW = 632.1 Da, observed adduct = 611.2 Da).

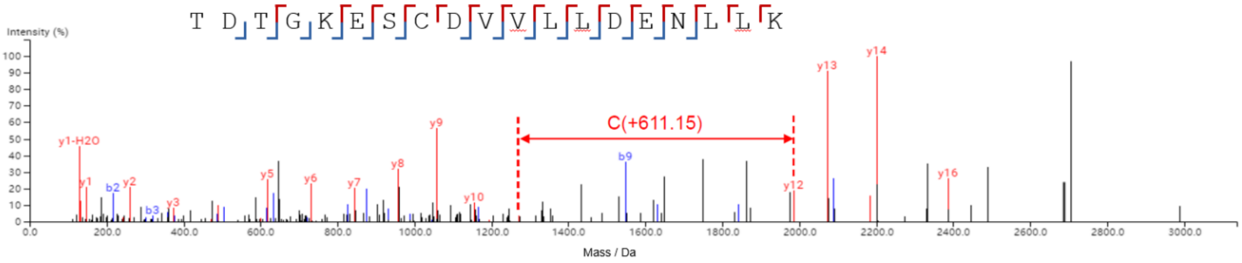

| # | b | b-H2O | b-NH3 | b(2+) | Seq | y | y-H2O | y-NH3 | y(2+) | # |
|---|---|---|---|---|---|---|---|---|---|---|
| 1 | 102.06 | 84.04 | 85.03 | 51.53 | T | | | | | 19 |
| 2 | 217.08 | 199.07 | 200.06 | 109.04 | D | 2602.15 | 2584.14 | 2585.12 | 1301.57 | 18 |
| 3 | 318.13 | 300.12 | 301.10 | 159.57 | T | 2487.12 | 2469.11 | 2470.10 | 1244.06 | 17 |
| 4 | 375.15 | 357.14 | 358.12 | 188.08 | G | 2386.07 | 2368.06 | 2369.05 | 1193.54 | 16 |
| 5 | 503.25 | 485.24 | 486.22 | 252.12 | K | 2329.05 | 2311.04 | 2312.03 | 1165.03 | 15 |
| 6 | 632.29 | 614.28 | 615.26 | 316.64 | E | 2200.96 | 2182.95 | 2183.95 | 1100.98 | 14 |
| 7 | 719.32 | 701.31 | 702.29 | 360.16 | S | 2071.92 | 2053.91 | 2054.89 | 1036.46 | 13 |
| 8 | 1433.48 | 1415.47 | 1416.45 | 717.24 | C(+611.15) | 1984.88 | 1966.87 | 1967.86 | 992.94 | 12 |
| 9 | 1548.51 | 1530.50 | 1531.48 | 774.75 | D | 1270.73 | 1252.71 | 1253.70 | 635.86 | 11 |
| 10 | 1647.58 | 1629.56 | 1630.56 | 824.28 | V | 1155.70 | 1137.69 | 1138.67 | 578.35 | 10 |
| 11 | 1746.64 | 1728.63 | 1729.62 | 873.83 | V | 1056.63 | 1038.62 | 1039.60 | 528.81 | 9 |
| 12 | 1859.73 | 1841.71 | 1842.70 | 930.38 | L | 957.56 | 939.55 | 940.53 | 479.28 | 8 |
| 13 | 1972.81 | 1954.80 | 1955.78 | 986.90 | L | 844.48 | 826.47 | 827.45 | 422.74 | 7 |
| 14 | 2087.86 | 2069.83 | 2070.81 | 1044.43 | D | 731.40 | 713.38 | 714.38 | 366.20 | 6 |
| 15 | 2216.88 | 2198.87 | 2199.85 | 1108.94 | E | 616.37 | 598.36 | 599.34 | 308.68 | 5 |
| 16 | 2330.92 | 2312.91 | 2313.90 | 1165.98 | N | 487.32 | 469.31 | 470.30 | 244.16 | 4 |
| 17 | 2444.01 | 2426.00 | 2426.98 | 1222.50 | L | 373.28 | 355.27 | 356.25 | 187.14 | 3 |
| 18 | 2557.09 | 2539.08 | 2540.07 | 1279.05 | L | 260.20 | 242.19 | 243.17 | 130.60 | 2 |
| 19 | | | | | K | 147.11 | 129.10 | 130.09 | 74.06 | 1 |

**Supplementary Figure 3. a:** Recombinant protein expression construct design for PIKfyve$^{WT}$ and PIKfyve$^{C1970A}$. **b,c:** SDS-PAGE analysis of purified recombinant PIKfyve$^{WT}$ (**b**) and PIKfyve$^{C1970A}$ (**c**), as visualized by Coomassie blue staining.

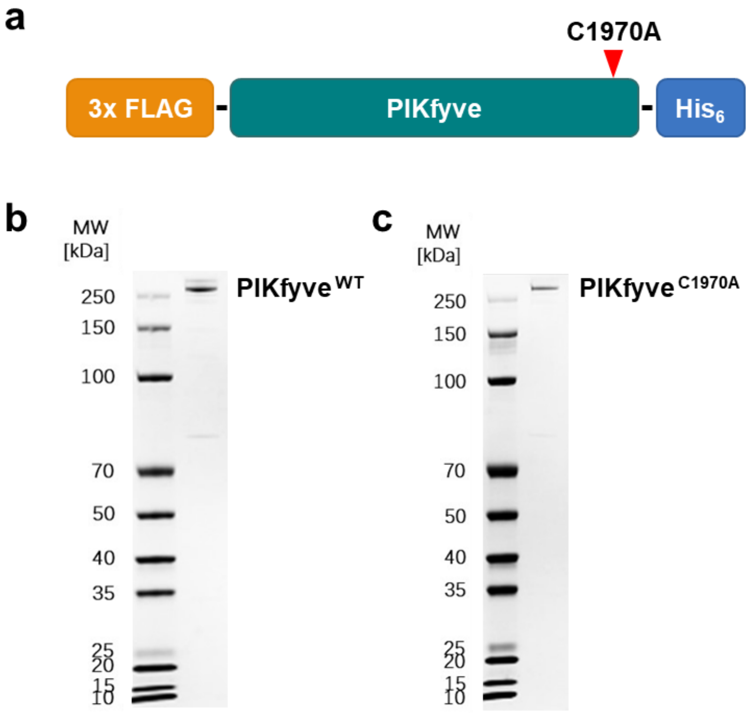

**Supplementary Figure 4. a:** The ATP $K_M^{app}$ of PIKfyve$^{WT}$ and PIKfyve$^{C1970A}$ indicate that the C1970A mutation does not affect kinase function. **b:** ADP-Glo assay results after 2 h of preincubation with reversible PIKfyve inhibitors YM-201636 (PIKfyve$^{WT}$ $IC_{50}$ = 23 nM (green circles), PIKfyve$^{C1970A}$ $IC_{50}$ = 26 nM (black squares)) and Staurosporine (PIKfyve$^{WT}$ $IC_{50}$ = 3.7 µM (green circles), PIKfyve$^{C1970A}$ $IC_{50}$ = 3.1 µM (black squares)). Both proteins are equally inhibited by the reversible inhibitors, indicating that C1970 does not influence their enzymatic activity.

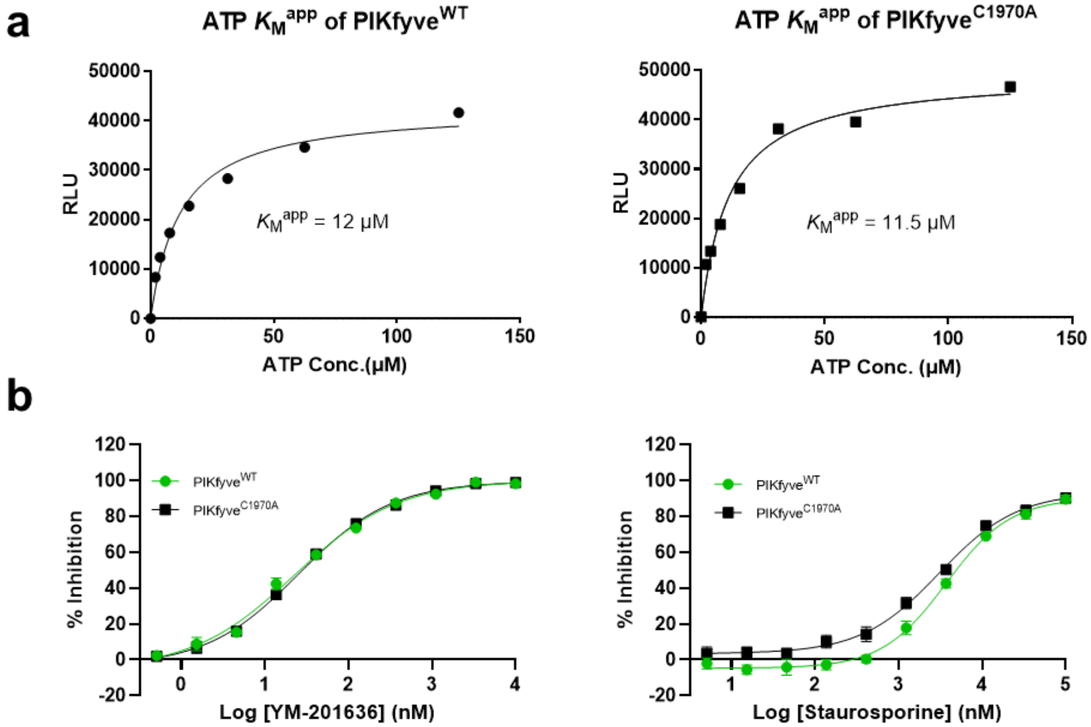

**Supplementary Figure 5. a:** SDS-PAGE analysis of PIKfyve$^{WT}$ treated overnight at 25 °C with Casp3, visualized by Coomassie blue staining. Selective liberation of the C-terminal 35 kDa kinase domain is achieved. * = HSP70 contaminant. **b:** Peptide mapping of the excised 35 kDa band after digestion with trypsin confirms the expected identity of the protein. Tryptic peptide coverage is highlighted in red.

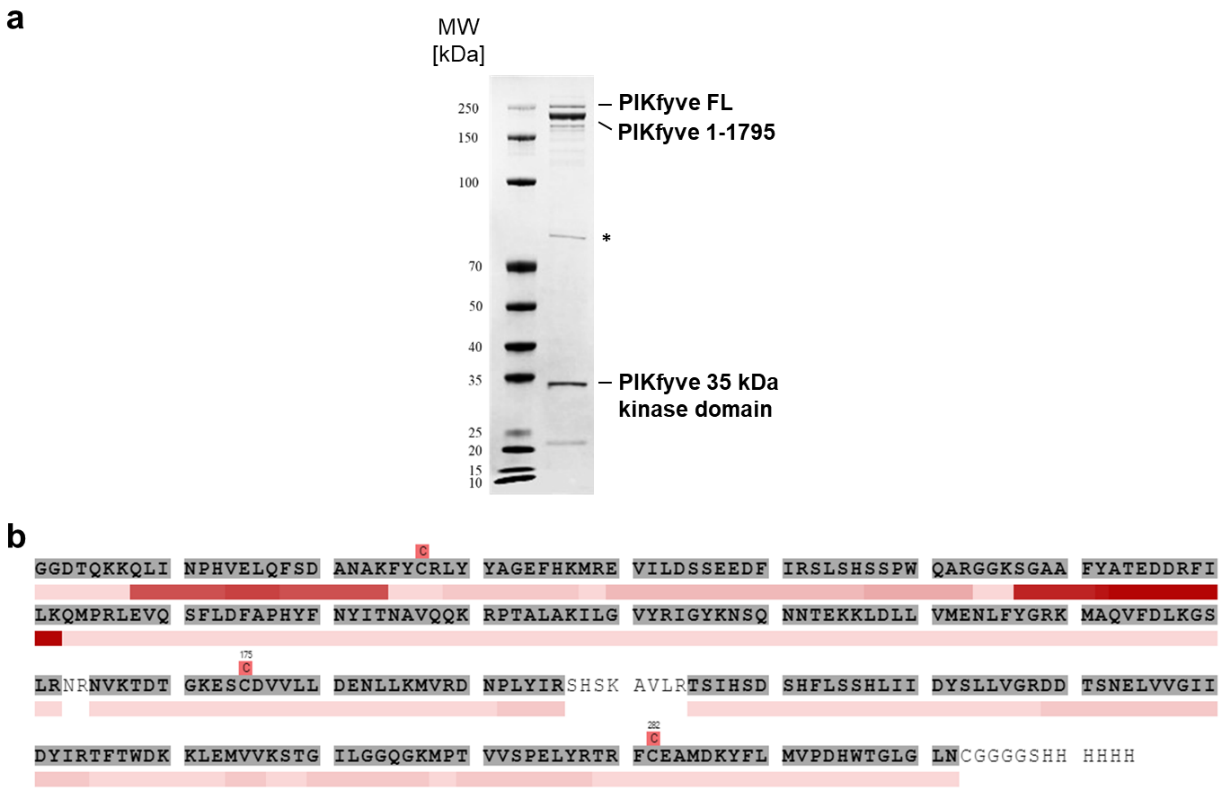

**Supplementary Figure 6.** Deconvoluted selective proteolysis-mediated intact mass analysis (IMA) spectrum for the reaction of DUN'465 with recombinant PIKfyve$^{WT}$ ([M+H]$^+$ = 36,525 Da (M+612 Da); [M+H]$^+$ = 37,137 Da (M+1,224 Da)). 82% single adduct and 18% double adduct are observed with PIKfyve$^{WT}$ protein after 1 h incubation at room temperature (0.42 μM PIKfyve$^{WT}$, 2.1 μM DUN'465).

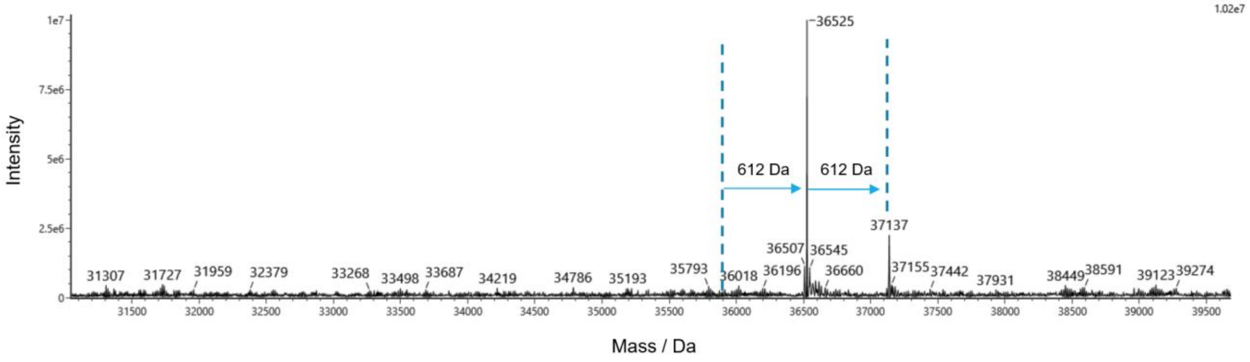

**Supplementary Figure 7.** LC-MS/MS analysis of trypsin-digested PIKfyve$^{WT}$ after treatment with DUN’058 (1 h, 25 °C) confirms covalent modification at C1970 (DUN’058 MW = 490.6 Da, observed adduct = 490.3 Da). No other modification sites were identified in the experiment (>90% sequence coverage observed).

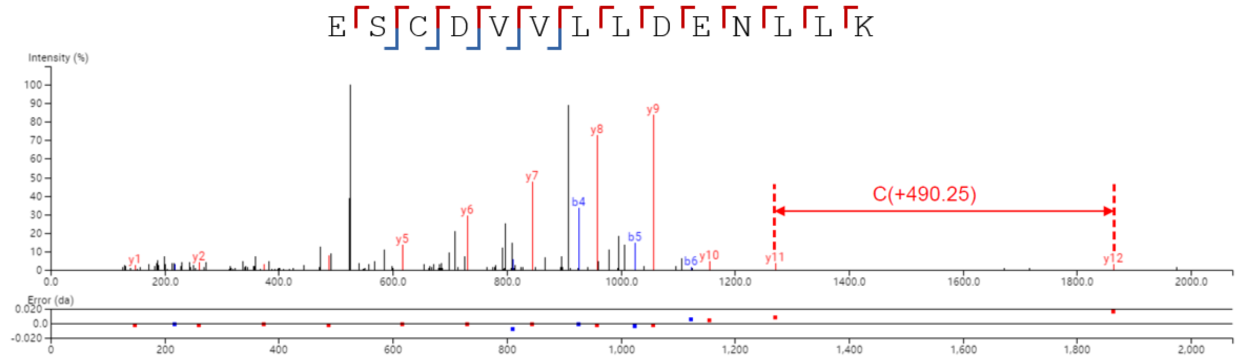

| # | b | b(2+) | Seq | y | y(2+) | # |
|---|---|---|---|---|---|---|
| 1 | 130.05 | 65.53 | E | | | 14 |
| 2 | 217.08 | 109.04 | S | 1951.02 | 976.01 | 13 |
| 3 | 810.35 | 405.67 | C(+490.25) | 1863.96 | 932.49 | 12 |
| 4 | 925.37 | 463.18 | D | 1270.71 | 635.86 | 11 |
| 5 | 1024.44 | 512.72 | V | 1155.69 | 578.35 | 10 |
| 6 | 1123.50 | 562.25 | V | 1056.63 | 528.81 | 9 |
| 7 | 1236.59 | 618.79 | L | 957.56 | 479.28 | 8 |
| 8 | 1349.67 | 675.34 | L | 844.48 | 422.74 | 7 |
| 9 | 1464.70 | 732.85 | D | 731.39 | 366.20 | 6 |
| 10 | 1593.74 | 797.37 | E | 616.37 | 308.68 | 5 |
| 11 | 1707.79 | 854.39 | N | 487.32 | 244.16 | 4 |
| 12 | 1820.87 | 910.93 | L | 373.28 | 187.14 | 3 |
| 13 | 1933.95 | 967.48 | L | 260.20 | 130.60 | 2 |
| 14 | | | K | 147.11 | 74.06 | 1 |

**Supplementary Figure 8.** Determination of $k_{inact}$ and $K_I$ for DUN'058 by selective-proteolysis mediated IMA. **a:** Occupancy of PIKfyve C1970 by DUN'058 is plotted as a function of time to generate an observed rate constant of modification for each concentration ($k_{obs}$). **b:** Plotting $k_{obs}$ as a function of DUN'058 concentration allows the fitting of $k_{inact}$ and $K_I$, and the subsequent calculation of $k_{inact}/K_I$.

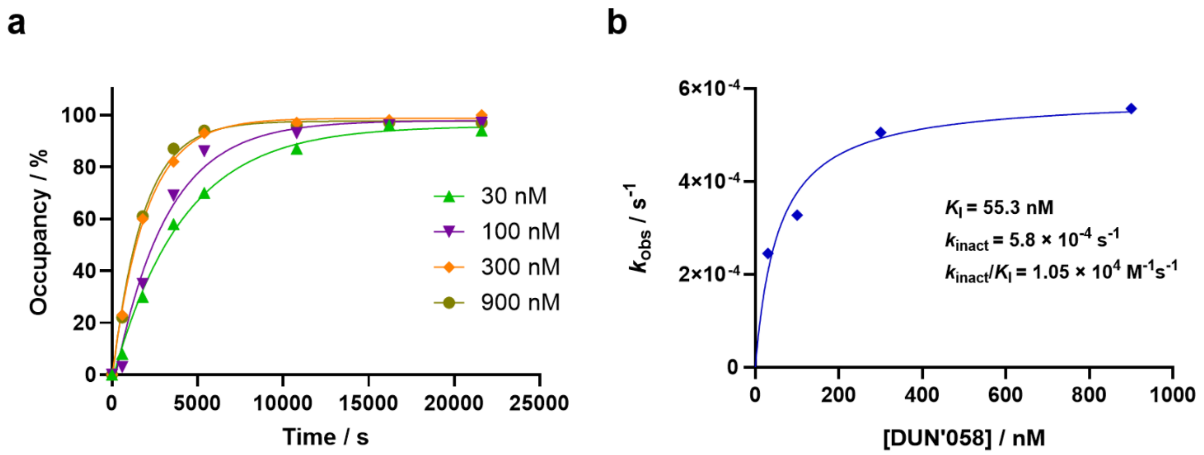

**Supplementary Figure 9. a:** Recombinant protein expression construct design and sequence for PIP4K2A$^{6mut}$, with mutations underlined and in bold. **b:** SDS-PAGE analysis of purified recombinant PIP4K2A$^{6mut}$, as visualized by Coomassie blue staining.

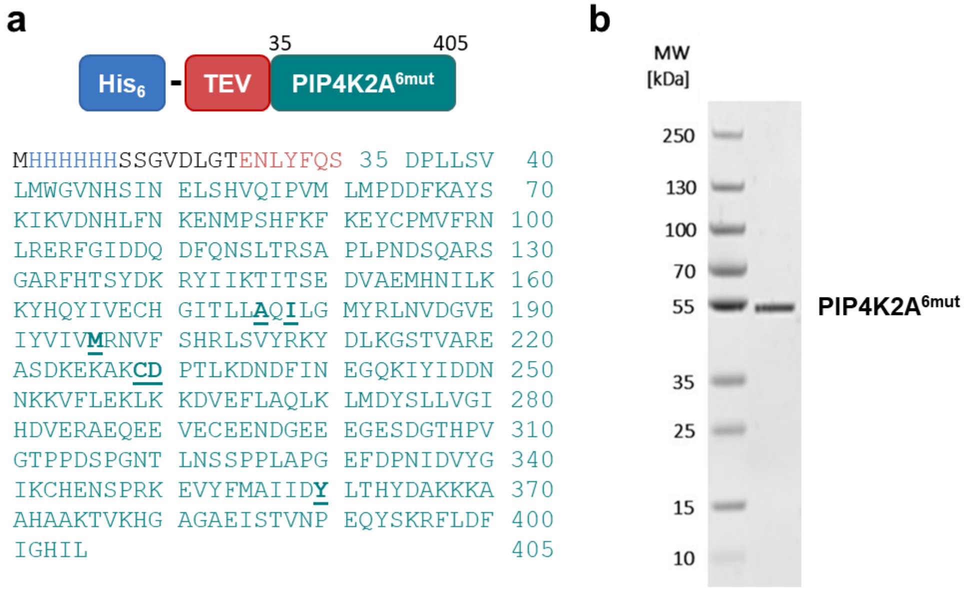

**Supplementary Figure 10.** Deconvoluted IMA spectrum for the treatment of 222 µM PIP4K2A$^{6mut}$ with 1 mM DUN’058 (6 h, 25 °C). A single adduct of DUN’058 with the protein was observed (DUN’058 MW = 490.6 Da; $[M+H]^+$ = 45,518 Da (M+489 Da)).

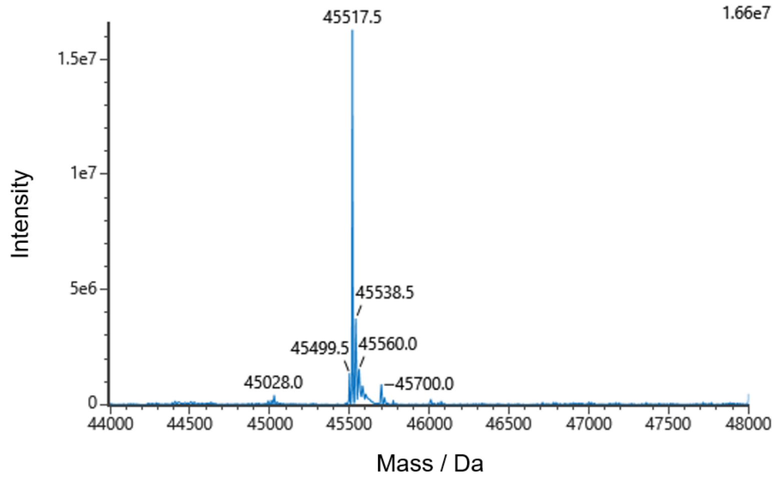

**Supplementary Figure 11.** LC-MS/MS analysis of trypsin-digested PIP4K2A$^{6mut}$ after treatment with DUN’058 (6 h, 25 °C) confirms covalent modification at C229 (DUN’058 MW = 490.6 Da, observed adduct = 490.3 Da). No other modification sites were identified in the experiment.

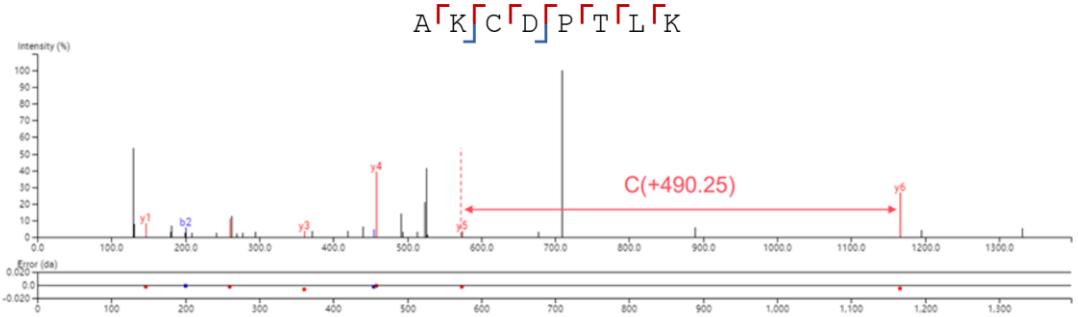

| # | b | b(2+) | Seq | y | y(2+) | # |
|---|---|---|---|---|---|---|
| 1 | 72.04 | 36.52 | A | | | 8 |
| 2 | 200.14 | 100.57 | K | 1294.68 | 647.84 | 7 |
| 3 | 793.40 | 397.20 | C(+490.25) | 1166.59 | 583.79 | 6 |
| 4 | 908.43 | 454.71 | D | 573.32 | 287.16 | 5 |
| 5 | 1005.48 | 503.24 | P | 458.30 | 229.65 | 4 |
| 6 | 1106.53 | 553.76 | T | 361.25 | 181.12 | 3 |
| 7 | 1219.61 | 610.30 | L | 260.20 | 130.60 | 2 |
| 8 | | | K | 147.11 | 74.06 | 1 |

**Supplementary Figure 12.** NanoBRET cellular target engagement assay data for DUN’058 and DUN’465. Treatment was performed at the indicated concentrations for 1 h in HEK293T cells. DUN’058 $IC_{50}$ = 3 nM (blue circles), DUN’465 $IC_{50}$ = 119 nM (grey triangles).

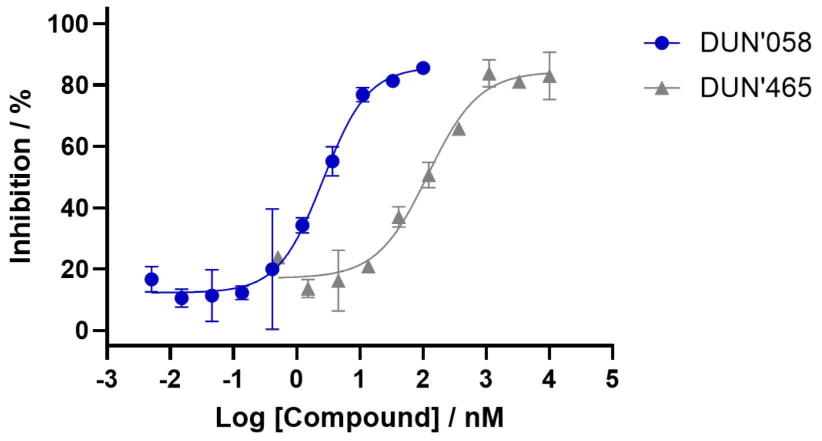

**Supplementary Figure 13.** Compound exposure (DUN'058) in the plasma (black circles) and brain (blue squares) of C57BL/6 mice after a single 50 mg/kg oral dose of DUN'058.

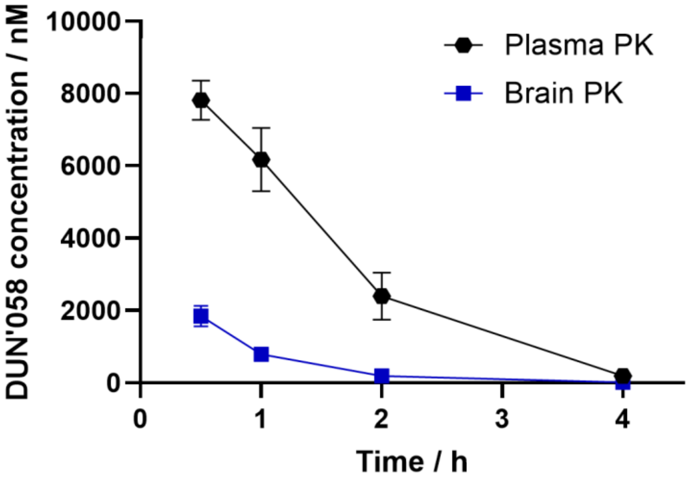

**Supplementary Figure 14.** The PIKfyve protein resynthesis rate was determined in brain and spleen by fitting TO data to a monoexponential decay model. The obtained first-order rate constants (*k*) in brain (**a**) and spleen (**b**) were converted to protein half-lives (brain = 72 h; spleen = 35 h). The PIKfyve half-life in duodenum was estimated to be < 24 h.

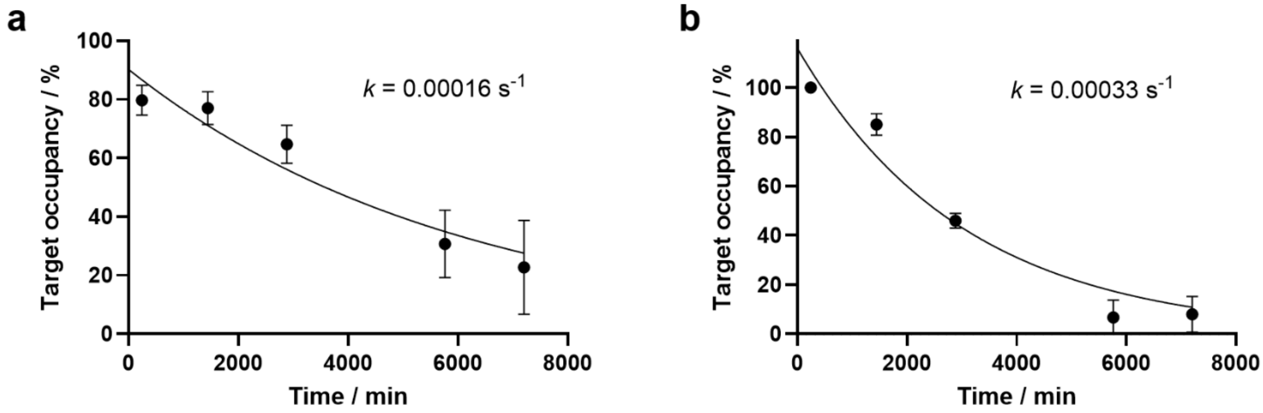

**Supplementary Figure 15.** 5-day *in vivo* repeat dose study data for DUN’058. Mice were treated daily with an oral dose of 50 mg/kg DUN’058 for 5 days. **a:** No change in body weight was observed on treatment with DUN’058 (green triangles) and vehicle (black circles). 0% body weight change is shown as a dotted line. **b:** Brain TO at PIKfyve C1970 for DUN’058, as determined by targeted MS, from tissue collected 0.5 and 24 h post last dose.

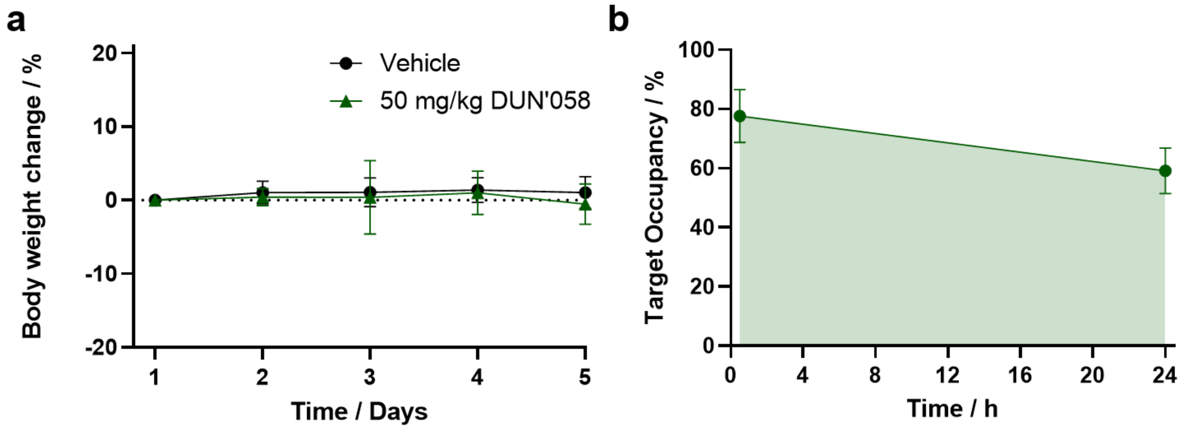

### 3. Supplementary Tables

**Supplementary Table 1:** *Summary of ABPP chemoproteomics data for PIKfyve C1970* can be found as an Excel document titled "SI_Table_1".

**Supplementary Table 2:** *DUN'465 ABPP data* can be found as an Excel document titled "SI_Table_2".

**Supplementary Table 4:** *DUN'058 ABPP data* can be found as an Excel document titled "SI_Table_4".

**Supplementary Table 6:** *DUN'058 global proteomics data* can be found as an Excel document titled "SI_Table_6".

**Supplementary Table 7:** *DUN'058 in vivo targeted MS data* can be found as an Excel document titled "SI_Table_7".

**Supplementary Table 3.** DUN’465 and DUN’058 kinome selectivity (measured at 1 μM) against a panel of 66 protein kinases and 4 lipid kinases, presented as percentage of enzyme activity.

| | % Enzyme Activity (Relative to DMSO Controls) | | | | % Enzyme Activity (Relative to DMSO Controls) | | | |
|---|---|---|---|---|---|---|---|---|
| Kinases | DUN'465 (1 μM) | | Control Compound $IC_{50}$ (M) | Control Compound ID | DUN'058 (1 μM) | | Control Compound $IC_{50}$ (M) | Control Compound ID |
| | Data 1 | Data 2 | | | Data 1 | Data 2 | | |
| ABL1 | 55.83 | 55.28 | 4.69E-08 | STAUROSPORINE | 89.39 | 88.92 | 3.02E-08 | STAUROSPORINE |
| AKT1 | 62.63 | 61.38 | 3.46E-09 | STAUROSPORINE | 101.46 | 99.34 | 3.66E-09 | STAUROSPORINE |
| ALK4/ACVR1B | 121.22 | 120.68 | 1.76E-07 | LDN193189 | 101.77 | 101.28 | 6.24E-08 | LDN193189 |
| ARK5/NUAK1 | 15.03 | 13.84 | 8.88E-10 | STAUROSPORINE | 96.04 | 95.83 | 6.30E-10 | STAUROSPORINE |
| Aurora A | 2.35 | 2.33 | 1.09E-09 | STAUROSPORINE | 64.64 | 64.63 | 1.10E-09 | STAUROSPORINE |
| BRAF | 102.19 | 97.78 | 1.17E-08 | GW5074 | 92.51 | 92.39 | 8.81E-09 | GW5074 |
| BTK | 1.46 | 1.37 | 1.83E-08 | STAUROSPORINE | 2.31 | 2.10 | 1.97E-08 | STAUROSPORINE |
| c-Kit | 11.65 | 11.42 | 1.16E-09 | STAUROSPORINE | 95.00 | 93.68 | 4.69E-10 | STAUROSPORINE |
| c-MET | 77.46 | 76.78 | 2.02E-08 | STAUROSPORINE | 104.41 | 103.12 | 2.17E-08 | STAUROSPORINE |
| c-Src | 18.44 | 17.99 | 2.63E-09 | STAUROSPORINE | 75.63 | 74.35 | 2.35E-09 | STAUROSPORINE |
| CAMK1a | 63.05 | 61.46 | 3.59E-09 | STAUROSPORINE | 96.39 | 96.09 | 2.78E-09 | STAUROSPORINE |
| CDK1/cyclin B | 59.66 | 59.07 | 1.26E-09 | STAUROSPORINE | 96.55 | 95.68 | 1.07E-09 | STAUROSPORINE |
| CDK4/cyclin D1 | 4.40 | 4.05 | 1.01E-08 | STAUROSPORINE | 92.19 | 91.59 | 8.31E-09 | STAUROSPORINE |
| CDK7/cyclin H | 32.74 | 32.62 | 5.74E-08 | STAUROSPORINE | 98.50 | 97.97 | 2.43E-08 | STAUROSPORINE |
| CDK9/cyclin T2 | 30.33 | 29.51 | 3.27E-09 | STAUROSPORINE | 95.76 | 95.45 | 7.01E-09 | STAUROSPORINE |
| CHK1 | 77.80 | 70.58 | 1.63E-10 | STAUROSPORINE | 94.95 | 94.72 | 1.36E-10 | STAUROSPORINE |
| CK1a1 | 102.64 | 101.48 | 5.05E-06 | STAUROSPORINE | 98.25 | 98.23 | 2.92E-06 | STAUROSPORINE |
| CK1g1 | 98.75 | 96.69 | 5.33E-06 | STAUROSPORINE | 106.21 | 102.71 | 1.08E-05 | STAUROSPORINE |
| CK2a | 75.69 | 75.67 | 1.64E-07 | GW5074 | 99.42 | 98.22 | 2.49E-07 | GW5074 |
| CLK2 | 8.36 | 8.14 | 1.67E-09 | STAUROSPORINE | 88.14 | 86.74 | 4.04E-09 | STAUROSPORINE |
| DAPK2 | 92.86 | 91.89 | 5.14E-09 | STAUROSPORINE | 97.63 | 97.52 | 6.06E-09 | STAUROSPORINE |
| DCAMKL1 | 45.28 | 44.18 | 6.16E-08 | STAUROSPORINE | 102.37 | 101.19 | 9.44E-08 | STAUROSPORINE |
| DYRK1/DYRK1A | 83.52 | 81.71 | 1.64E-09 | STAUROSPORINE | 88.00 | 87.22 | 2.39E-09 | STAUROSPORINE |
| EGFR | 3.66 | 3.65 | 7.10E-08 | STAUROSPORINE | 56.08 | 55.77 | 1.31E-07 | STAUROSPORINE |
| EPHA5 | 93.14 | 91.48 | 1.26E-08 | STAUROSPORINE | 99.39 | 98.66 | 1.75E-08 | STAUROSPORINE |
| EPHB2 | 106.63 | 106.62 | 6.66E-08 | STAUROSPORINE | 104.37 | 104.33 | 7.25E-08 | STAUROSPORINE |
| ERK1 | 96.29 | 95.97 | 5.53E-09 | SCH772984 | 93.77 | 91.54 | 1.76E-09 | SCH772984 |
| FGFR2 | 1.30 | 1.23 | 2.98E-09 | STAUROSPORINE | 59.83 | 57.80 | 1.65E-09 | STAUROSPORINE |
| FLT1/VEGFR1 | 49.15 | 46.16 | 5.91E-09 | STAUROSPORINE | 98.47 | 96.27 | 4.63E-09 | STAUROSPORINE |
| FLT3 | 1.12 | 0.75 | 6.50E-10 | STAUROSPORINE | 82.90 | 82.53 | 1.62E-09 | STAUROSPORINE |
| GSK3b | 84.57 | 83.84 | 4.26E-09 | STAUROSPORINE | 93.39 | 93.26 | 5.23E-09 | STAUROSPORINE |
| HIPK2 | 5.20 | 5.11 | 1.07E-07 | STAUROSPORINE | 94.88 | 94.51 | 2.55E-07 | STAUROSPORINE |
| IGF1R | 103.45 | 101.92 | 3.93E-08 | STAUROSPORINE | 95.51 | 94.45 | 2.69E-08 | STAUROSPORINE |
| IKKe/IKBKE | 66.29 | 63.51 | 2.09E-10 | STAUROSPORINE | 95.57 | 95.49 | 2.74E-10 | STAUROSPORINE |
| IRAK4 | 57.87 | 57.45 | 6.38E-09 | STAUROSPORINE | 93.04 | 92.02 | 7.07E-09 | STAUROSPORINE |
| JAK2 | 2.41 | 2.35 | 1.59E-10 | STAUROSPORINE | 94.71 | 93.87 | 1.99E-10 | STAUROSPORINE |
| JNK3 | 5.21 | 5.14 | 4.37E-08 | JNKI VIII | 89.70 | 89.31 | 3.79E-08 | JNKI VIII |
| LIMK1 | 11.01 | 10.96 | 5.66E-10 | STAUROSPORINE | 3.47 | 3.46 | 4.91E-10 | STAUROSPORINE |
| LOK/STK10 | 57.81 | 56.99 | 5.30E-08 | RO-31-8220 | 91.83 | 89.71 | 1.66E-08 | RO-31-8220 |
| LYN | 23.68 | 23.45 | 7.70E-10 | STAUROSPORINE | 96.15 | 94.44 | 8.58E-10 | STAUROSPORINE |
| MARK2/PAR-1Ba | 61.85 | 60.46 | 9.81E-11 | STAUROSPORINE | 97.65 | 96.42 | 6.51E-11 | STAUROSPORINE |
| MAST3 | 99.23 | 98.30 | 1.08E-06 | STAUROSPORINE | 99.93 | 98.95 | 7.64E-07 | STAUROSPORINE |
| MEK1 | 96.88 | 96.68 | 1.32E-08 | STAUROSPORINE | 87.93 | 86.06 | 1.66E-08 | STAUROSPORINE |
| MEKK1 | 100.61 | 98.31 | 5.69E-07 | STAUROSPORINE | 100.35 | 100.12 | 1.14E-06 | STAUROSPORINE |
| MLK1/MAP3K9 | 4.60 | 4.53 | 1.21E-09 | STAUROSPORINE | 91.08 | 90.62 | 1.03E-09 | STAUROSPORINE |
| MNK2 | 65.54 | 64.43 | 4.33E-08 | STAUROSPORINE | 96.04 | 96.02 | 2.41E-08 | STAUROSPORINE |
| MSK2/RPS6KA4 | 78.40 | 76.88 | 1.79E-09 | STAUROSPORINE | 100.95 | 98.57 | 1.55E-09 | STAUROSPORINE |
| MST1/STK4 | 103.10 | 102.35 | 1.64E-09 | STAUROSPORINE | 97.40 | 95.78 | 1.25E-09 | STAUROSPORINE |
| MYLK3 | 99.57 | 97.24 | 5.99E-08 | STAUROSPORINE | 99.47 | 98.87 | 8.47E-08 | STAUROSPORINE |
| NEK1 | 1.68 | 1.23 | 1.36E-08 | STAUROSPORINE | 96.19 | 88.81 | 2.92E-08 | STAUROSPORINE |
| P38a/MAPK14 | 110.03 | 105.53 | 1.65E-08 | SB202190 | 98.54 | 98.09 | 1.48E-08 | SB202190 |
| p70S6K/RPS6KB1 | 11.55 | 11.07 | 4.72E-10 | STAUROSPORINE | 87.00 | 84.68 | 4.41E-10 | STAUROSPORINE |
| PAK4 | 89.61 | 85.89 | 3.09E-09 | STAUROSPORINE | 97.84 | 95.34 | 2.98E-09 | STAUROSPORINE |
| PIM3 | 96.04 | 94.13 | 9.93E-11 | STAUROSPORINE | 73.23 | 72.38 | 6.43E-11 | STAUROSPORINE |
| PKA | 52.03 | 51.96 | 1.64E-09 | STAUROSPORINE | 91.47 | 89.86 | 1.63E-09 | STAUROSPORINE |
| PKCa | 69.67 | 69.17 | 1.99E-10 | STAUROSPORINE | 91.52 | 89.85 | 1.38E-10 | STAUROSPORINE |
| PKCtheta | 79.92 | 79.30 | 1.83E-10 | STAUROSPORINE | 84.15 | 80.51 | 1.65E-10 | STAUROSPORINE |
| PLK1 | 89.39 | 89.00 | 1.42E-07 | STAUROSPORINE | 101.05 | 99.03 | 1.71E-07 | STAUROSPORINE |
| RET | 1.55 | 1.51 | 2.88E-09 | STAUROSPORINE | 90.49 | 89.87 | 5.24E-09 | STAUROSPORINE |
| ROCK1 | 81.63 | 80.38 | 7.00E-10 | STAUROSPORINE | 99.27 | 96.95 | 9.85E-10 | STAUROSPORINE |
| RSK1 | 75.11 | 71.66 | 1.24E-10 | STAUROSPORINE | 91.79 | 89.63 | 1.25E-10 | STAUROSPORINE |
| STK22D/TSSK1 | 34.78 | 34.27 | 3.11E-11 | STAUROSPORINE | 94.42 | 94.07 | 4.21E-11 | STAUROSPORINE |
| TEC | 15.77 | 15.75 | 5.01E-08 | STAUROSPORINE | 17.13 | 16.34 | 4.56E-08 | STAUROSPORINE |
| TGFBR2 | 94.37 | 93.15 | 1.19E-07 | LDN193189 | 97.54 | 97.32 | 5.71E-08 | LDN193189 |
| TRKA | 36.49 | 35.97 | 2.00E-09 | STAUROSPORINE | 90.99 | 90.27 | 1.81E-09 | STAUROSPORINE |
| ULK1 | 3.22 | 3.06 | 7.81E-09 | STAUROSPORINE | 100.02 | 98.50 | 8.55E-09 | STAUROSPORINE |
| PI3Ka (p110a/p85a) | 93.40 | 92.87 | 4.00E-09 | PI-103 | 116.50 | 116.46 | 8.29E-09 | PI-103 |
| PI3Kb (p110b/p85a) | 96.07 | 95.42 | 1.33E-08 | PI-103 | 152.75 | 140.74 | 3.35E-08 | PI-103 |
| PI3Kd (p110d/p85a) | 96.65 | 96.04 | 3.60E-09 | PI-103 | 83.15 | 82.47 | 1.45E-08 | PI-103 |
| PI3Kg (p110g) | 100.11 | 96.19 | 2.31E-08 | PI-103 | 111.18 | 106.80 | 5.61E-08 | PI-103 |

**Supplementary Table 5.** Data collection and refinement statistics for X-ray structure of PIP4K2A$^{6mut}$ bound to DUN'058

| | PIP4K2A$^{6mut}$ bound to DUN'058 |
|---|---|
| **PDB ID** | 9zmo |
| **Data collection** | |
| Space group | $P4_22_12$ |
| Cell dimensions | |
| *a*, *b*, *c* (Å) | 112.26, 112.26, 192.80 |
| α, β, γ (°) | 90.00, 90.00, 90.00 |
| Resolution (Å) | 97.01-3.08 (3.39-3.08)* |
| $R_{sym}$ or $R_{merge}$ | 0.469 (2.94) |
| $CC_{1/2}$ | 0.998 (0.616) |
| $I / \sigma I$ | 7.1 (1.4) |
| Completeness (%) | 94.6 (59.8) |
| Redundancy | 20.1 |
| **Refinement** | |
| Resolution (Å) | 3.08 |
| No. reflections | 15,420 |
| $R_{work}$ / $R_{free}$ | 0.210 / 0.270 |
| No. atoms | 5,112 |
| Protein | 5,037 |
| Ligand/ion | 75 |
| *B*-factors | |
| Mean *B*-factor (Å$^2$) | 97.02 |
| R.m.s. deviations | |
| Bond lengths (Å) | 0.007 |
| Bond angles (°) | 1.73 |

*Values in parentheses refer to the highest resolution shell (3.39-3.08 Å).
